# Intestinal serotonergic vagal signaling as a mediator of microbiota-induced hypertension

**DOI:** 10.1101/2024.07.17.603451

**Authors:** Alan de Araujo, Hemaa Sree Kumar, Tao Yang, Adriana Alviter Plata, Elliott W. Dirr, Nicole Bearss, David M. Baekey, Darren S. Miller, Basak Donertas-Ayaz, Niousha Ahmari, Arashdeep Singh, Andrea L. Kalinoski, Timothy J. Garrett, Christopher J. Martyniuk, Guillaume de Lartigue, Jasenka Zubcevic

**Affiliations:** Monell Chemical Senses Center, University of Pennsylvania; Department of Physiology and Pharmacology, University of Toledo; Department of Neuroscience, University of Florida; Department of Physiological Sciences, University of Florida; Department of Surgery, University of Toledo; Department of Pathology, Immunology and Laboratory Medicine, University of Florida.

**Keywords:** serotonin, vagus, GI, SHR, microbiota, hypertension, FMT, blood pressure, 5ht3a receptors

## Abstract

Hypertension is a pervasive global health challenge, impacting over a billion individuals worldwide. Despite strides in therapeutic strategies, a significant proportion of patients remain resistant to the currently available therapies. While conventional treatments predominantly focus on cardiac, renal, and cerebral targets, emerging research underscores the pivotal role of the gut and its microbiota. Yet, the precise mechanisms governing interactions between the gut microbiota and the host blood pressure remain unclear. Here we describe a neural host-microbiota interaction that is mediated by the intestinal serotonin (5-HT) signaling via vagal 5HT3a receptors and which is crucial for maintenance of blood pressure homeostasis. Notably, a marked decrease in both intestinal 5-HT and vagal 5HT3aR signaling is observed in hypertensive rats, and in rats subjected to fecal microbiota transplantation from hypertensive rats. Leveraging an intersectional genetic strategy in a Cre rat line, we demonstrate that intestinal 5HT3aR vagal signaling is a crucial link between the gut microbiota and blood pressure homeostasis and that recovery of 5-HT signaling in colon innervating vagal neurons can alleviate hypertension. This paradigm-shifting finding enhances our comprehension of hypertensive pathophysiology and unveils a promising new therapeutic target for combating resistant hypertension associated with gut dysbiosis.

## Introduction

Blood pressure is of fundamental importance for the fitness of the whole organism, and chronically elevated blood pressure or hypertension, when left untreated, invariably leads to cardiovascular, renal, and neurological conditions [1–3]. Despite the increasing prevalence of hypertension affecting over 1.5 billion people globally, its multiple etiologies remain a therapeutic challenge as a significant proportion of patients experience resistant hypertension [4–6], underlining the need for novel therapeutic strategies.

Recent research points to the role of the gut microbiota in the etiology of hypertension. The gastrointestinal (GI) tract harbors a vast and complex ecosystem of microbes that influence various physiological processes including blood pressure. Several studies reveal a link between gut dysbiosis, a term that describes an imbalance in gut microbes, and high blood pressure in rodents and human patients [7–9] but the host-microbiota interactions in the GI tract that maintain blood pressure homeostasis remain unclear.

Here, we describe a new mechanism of host-microbiota interaction mediated by intestinal serotonin (5-HT) signaling that is critical for maintenance of blood pressure homeostasis. We identified a genetically defined population of intestinal serotonergic vagal neurons that respond to changes in the gut microbiota and whose signaling is reduced in dysbiosis-induced hypertension. We show that these neurons are critical for blood pressure homeostasis and that recovery of intestinal vagal 5-HT signaling can mitigate dysbiosis-induced hypertension. These results reveal new evidence for the causal role of gut vagal afferents in regulation of blood pressure and highlight the therapeutic potential of recruiting the host-microbiota axis to reduce the burden of hypertension.

## Results

### Fecal transplant of hypertensive microbiota reduces colonic serotonin and elevates blood pressure

Several studies have linked gut dysbiosis to hypertension [7, 10], but causative mechanisms are unclear. To identify these, we employed fecal microbiota transplantation (FMT) in normotensive Wistar Kyoto rats (WKY). To deplete the endogenous gut bacteria, male WKY rats received a five-day course of oral antibiotics [11], resulting in approximately 75% decrease in fecal biomass (Extended data Fig. 1A). We then transplanted the whole gut microbiota slurry from either the spontaneously hypertensive rats (SHR) or their normotensive genetic controls, the WKY rats, into the antibiotic-depleted recipient WKY rats (Fig. 1A) and confirmed by the 16S rRNA gene sequencing at endpoint (Fig. 1B). In our model, the gut bacterial communities of FMT recipients showed compositional similarity to those of their respective donors, as illustrated by the box and principal component plots highlighting alpha and beta diversity (Fig. 1B). The microbiota differences between the two FMT groups were further highlighted by reduced fecal butyrate in the WKY recipients of SHR microbiota (i.e., the SHRèWKY group) compared to the recipients of the WKY microbiota (the control WKYèWKY group) (Fig 1B) consistent with our previous report of reduced bacterial butyrate producers in hypertension [7]. Telemetric recordings demonstrated a significant time-dependent rise in blood pressure in the SHRèWKY group compared to the WKY→WKY controls, peaking at week 8 post-FMT, with no major effects on the heart rate (Fig. 1D and Extended Data Fig. 1E), in agreement with previous studies that demonstrated a link between gut dysbiosis and hypertension in rodents [12, 13].

**Fig. 1:**
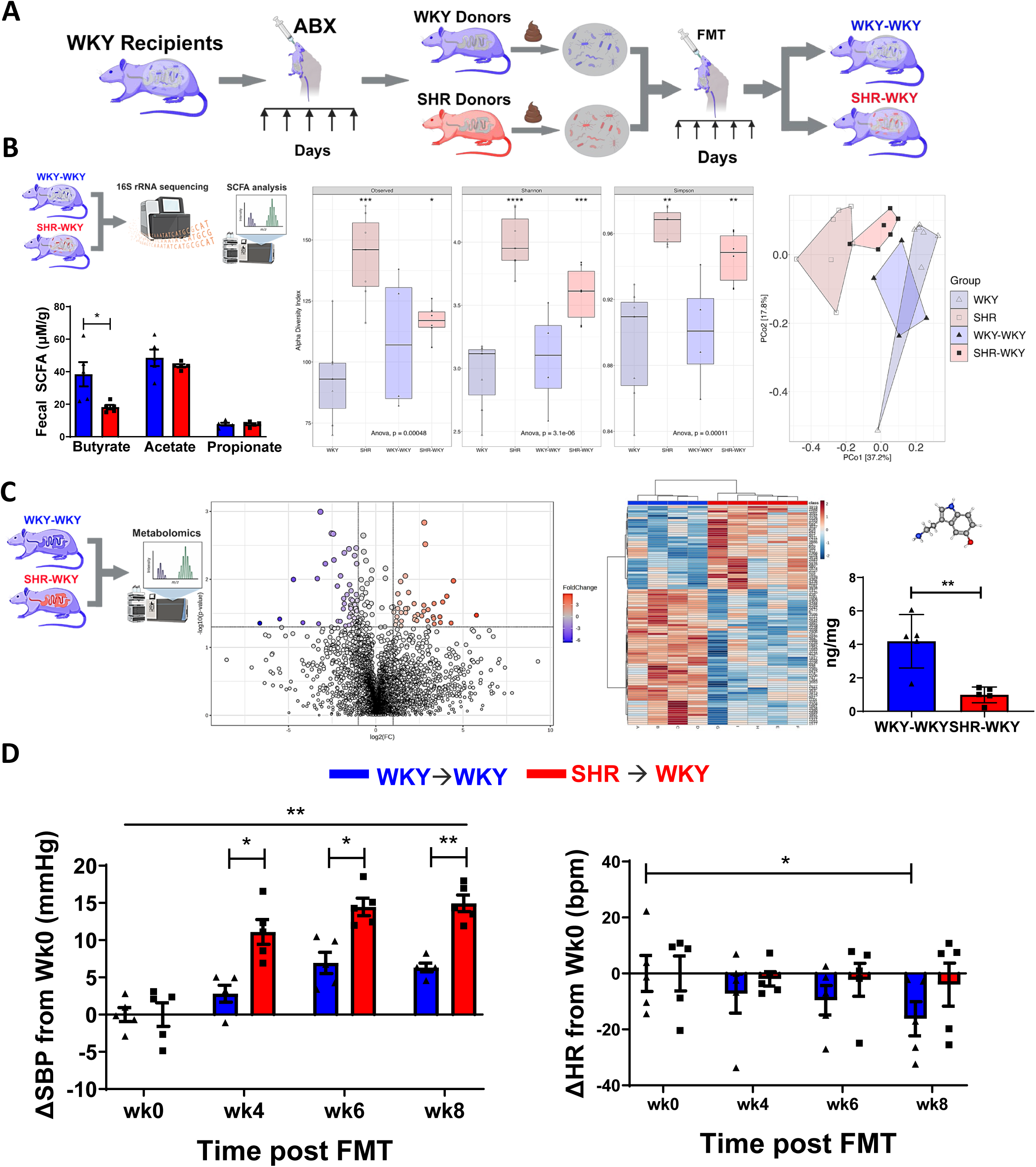
Transplant of hypertensive microbiota increased blood pressure and reduced colonic serotonin. **A,** cartoon schematics (created in Biorender) illustrating fecal microbiota transplant (FMT). Normotensive Wistar-Kyoto (WKY) rats were administered a cocktail of antibiotics (ABX) by daily oral gavage for five days, followed by oral gavage of fecal slurry from either the normotensive WKY or the spontaneously hypertensive rat (SHR) donors for five consecutive days, then once a week for eight weeks (1:1 donor to recipient ratio). Two groups were generated: WKY→WKY (blue) and SHR→WKY (red) reflecting donor→recipient microbiota transplant. **B,** top, schematics of endpoint 16S bacterial sequencing performed in all donor and recipient groups (N=4-7/group), and short chain fatty acid (SCFA) analyses performed in fecal samples of the two FMT recipient groups (N=5/group). Below, FMT from SHR to WKY produced a ∼2fold reduction in fecal butyrate (red bars). In the middle and far right panels, alpha and beta diversity plots following 16S bacterial sequencing analyses show bacterial abundance and diversity highlighting statistical similarities between the WKY and SHR donors and their respective gut bacteria recipient groups. Values are means ± SEM; ANOVA with Mann-Whitney post-hoc, *P<0.05, **P<0.01, ***P<0.001. **C**, on the left, cartoon schematics (created in Biorender) of endpoint metabolomics performed in proximal colons of WKY→WKY (blue) and SHR→WKY (red) rats using LC-MS. The two middle panels reflect significant divergence of overall metabolites between the WKY→WKY (blue) and SHR→WKY (red) reflected in the volcano plot and heat map (N=4-5/group). Far right panel illustrates a ∼4 fold decrease in serotonin (5-HT) in the proximal colons of WKY→WKY (blue) compared to the SHR→WKY (red) (N=5/group). Values are means ± SEM; Student’s Ttest, **P<0.01. **D**, Radiotelemetric measurements of continuous systolic blood pressure (SBP, mmHg) and heart rate (HR, beats per minute - bpm) were performed once a week over 24hrs in conscious unrestrained WKY→WKY (blue) and SHR→WKY (red) rats at baseline (i.e., before fecal microbiota transplant (FMT), noted as wk0) and then at weeks 4-8. In the left panel, a significant increase in SBP (by ∼10mmHg) was observed in the SHR→WKY group (red) compared to WKY→WKY control group (blue) starting at wk4 post FMT (N=5/group). Line across all time points reflects a significant interaction between treatment and time. In the right panel, no major differences were observed in HR between the groups, apart from a significant decrease in HR (by ∼20 bpm) that was observed at wk8 in the WKY→WKY control group (blue) compared to baseline (wk0). To better illustrate change from baseline for each rat and group, weekly absolute SBP and HR values were normalized to values observed at baseline (wk0) for each rat and presented as change (Δ) from wk0. Values are means ± SEM; ANOVA with Mann-Whitney post-hoc, *P<0.05, **P<0.01.

The proximal colon is the principal site of host-microbiota interactions mediated in part via SCFAs such as butyrate [14]. Thus, we performed metabolomic analyses of the whole proximal colon samples from WKY→WKY and SHR→WKY recipient groups and their respective donors using high-resolution liquid chromatography coupled with mass spectrometry (LC–MS/MS, Fig. 1C). Altered abundances of overall colonic metabolites were noted between the two FMT groups (Fig. 1C), indicating a broader effect of hypertensive microbiota on the colonic environment. Previous studies reported deregulations in tryptophan (Trp) metabolism in human and rodent hypertension [15–17]. The GI tract contains most of the total body 5-HT [18, 19], and our analyses of colonic metabolites in FMT rats identified several Trp metabolites. However, the only significant difference observed was a ∼4-fold reduction in 5-HT in the colons of SHR→WKY compared to WKY→WKY (Fig. 1C), mirroring what we observed in the colons of WKY and SHR donors (Extended Data Fig. 1C). As no difference was observed in fecal Trp or 5-HT in WKY→WKY and SHR→WKY rats (Extended Data Fig. 1D and Extended Data Table 6), this demonstrated no direct contribution of bacterial-derived Trp to the observed reduction in host colonic 5-HT in the SHR→WKY rats. Notably, FMT from SHR donors reduced relative expression of colonic tryptophan hydroxylase 1 (Tph1) in WKY recipients (Extended data Fig. 1F), a rate-limiting enzyme responsible for conversion of Trp to 5-HT in the intestinal enteroendocrine cells [19]. These data demonstrate a novel finding that colonic 5-HT dysfunction can be transplanted with the hypertensive microbiota and align with previous reports linking hypertension with deregulated Trp metabolism and 5-HT signaling [15–17, 20].

### A pivotal role for vagal serotonin signaling in blood pressure homeostasis

5-HT has known effects on vascular beds which can modulate blood pressure [15], but whether intestinal 5-HT can directly regulate blood pressure was unknown. Intestinal 5-HT signaling is in part mediated via the 5-HT receptor 3a (5HT3aR) expressed on vagal sensory neurons projecting to the nodose ganglia (NG^5ht3aR^) [21–25]. We first investigated the role of NG^5ht3aR^neurons in blood pressure homeostasis by employing optogenetics to selectively stimulate NG^5ht3aR^ neurons and measure the effects on blood pressure and heart rate. For this, we expressed the light-sensitive ion channel channelrhodopsin-2 (Chr2) in the NG^5ht3aR^ neurons of LE-5HT3a*^em1(T2A-Cre)Sage^* (5HT3aR^Cre^) rats (Fig. 2A). Bilateral NG microinjections of Cre-dependent AAV were used to deliver ChR2 (AAV-DIO-ChR2) in the NG^5ht3aR^neurons, while control rats received an AAV not carrying ChR2 (AAV-DIO-tdTomato, see Extended Data for details on vectors) (Fig. 2A and 2D). We first confirmed the model by observing virally mediated gene expression in the NG, the nucleus of the solitary tract (NTS), and the proximal colons of injected 5HT3aR^Cre^ rats (Fig. 2D and Extended Data Fig. 2A). We then bilaterally stimulated NG^5ht3aR^ neurons in anaesthetized rats by shining the excitatory blue light on the NG [26–28]. This triggered an immediate and time-locked increase in vagal afferent firing (Fig. 2B) and resulted in a pronounced decrease in blood pressure (∼20 mmHg, Fig. 2C and 2E) and a reduction in heart rate (by ∼40 beats per minute, bpm, Fig. 2C and 2F) in rats expressing the ChR2. The effects typically persisted for the entire duration of light stimulation and slowly recovered to baseline following cessation of the stimulus (Fig. 2C). In contrast, rats injected with control virus exhibited no significant changes in vagal firing, blood pressure or heart rate in response to optic stimulation of NG^5ht3aR^ neurons (Fig. 2A-F). These findings revealed a previously unknown role for NG^5ht3aR^ signaling in short-term regulation of blood pressure and heart rate.

**Fig. 2:**
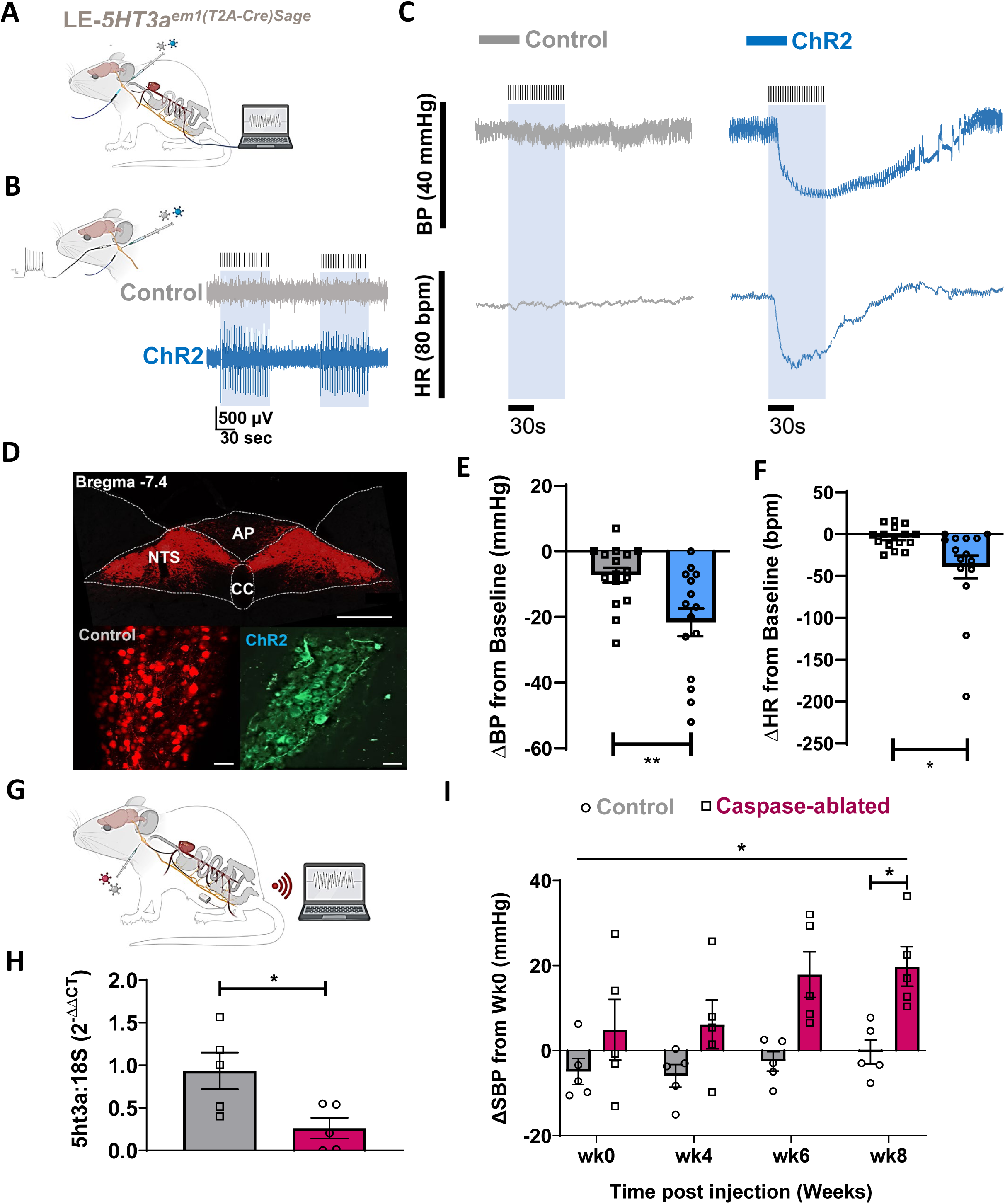
Vagal afferents sensing serotonin can modify blood pressure. **A,** Cre-dependent viral vectors (AAV) expressing either excitatory Channelrhodopsin 2 (ChR2) or control tdTomato (Control) were bilaterally microinjected in the nodose ganglia of LE-5HT3a^em1(T2A-Cre)Sage^ (5HT3aR^Cre^) rats. Three weeks later, rats were anaesthetized, and femoral artery was cannulated for measurement of direct blood pressure (BP, mmHg) and heart rate (HR, beats per minute, bpm) before, during and after application of the blue excitatory light to nodose ganglia for one minute. Schematics created in Biorender. **B,** an example of optic stimulation of the nodose ganglia expressing the excitatory ChR2 (in blue) and the control tdTomato (Control, in gray). Blue light stimulation produced a time-locked activation of vagal afferents expressing only the excitatory ChR2 in 5HT3aR^Cre^ rats. Light blue box illustrates timing of optic stimulation. **C**, representative traces of BP and HR before, during and after optic stimulation of the nodose ganglia expressing the excitatory ChR2 or control tdTomato. Optic stimulation of nodose ganglia expressing ChR2 produced a time-locked decrease in BP and HR (in blue) while optic stimulation of the nodose ganglia expressing the control tdTomato (in gray) did not produce a significant response in 5HT3aR^Cre^ rats. Light blue box illustrates timing of optic stimulation. **D**, endpoint confirmation of viral expression of Control (tdTomato, in red) in the nucleus of the solitary tract (NTS, top panel, scale bar=500µm) and nodose ganglia (bottom left), and ChR2 (in green) in the nodose ganglia (bottom right) (scale bar=100 µm). AP-area postrema; CC-central canal. **E**, average BP (mmHg) responses to bilateral optic stimulation of the nodose ganglia expressing Chr2 (blue bar) and Control (tdTomato, gray bar) (N=8 rats per group). Absolute BP values are normalized to values observed at baseline for each rat and presented as change (Δ) from baseline. Values are means ± SEM; Student’s Ttest, **P<0.01. **F**, average HR (bpm) responses to bilateral optic stimulation of the nodose ganglia expressing Chr2 (blue bar) and Control (tdTomato, gray bar) (N=8 rats per group). Absolute HR values are normalized to values observed at baseline for each rat and presented as change (Δ) from baseline. Values are means ± SEM; Student’s Ttest, *P<0.05. **G,** radiotelemetry transmitters were implanted in femoral artery of 5HT3aR^Cre^ rats for continuous measurement of BP and HR in conscious unrestrained rats. Following baseline measurements of BP and HR (wk0), Cre-dependent viral vectors (AAV) expressing either Caspase 3 (Cas, magenta), to selectively ablate 5HT3aR-expressing vagal afferents, or control tdTomato (gray) were bilaterally microinjected in the nodose ganglia in separate 5HT3aR^Cre^ rats. **H,** at endpoint, virally mediated ablation of 5HT3aR-expressing vagal afferent neurons was confirmed by reduced relative expression levels of nodose ganglia 5HT3aRs in Cas-injected rats (magenta) as measured by quantitative real time PCR (qRT-PCR). N=5/group. Values are means ± SEM; Student’s Ttest, *P<0.05. **I**, radiotelemetric measurements of continuous systolic BP (SBP, mmHg) were performed once a week over 24hrs in conscious unrestrained 5HT3aR^Cre^ rats at baseline (i.e., before nodose ganglia viral injections, noted as wk0) and then at weeks 4-8. A significant increase in SBP (by∼18mmHg) was observed in the Cas-injected group (magenta) compared to Control group (gray) at wk8 post injection (N=5/group). Line across all time points reflects significant interaction between treatment and time. Absolute values are normalized to values observed at baseline for each rat and presented as change (Δ) from baseline. Values are means ± SEM; ANOVA with Mann-Whitney post-hoc, *P<0.05.

Hypertension is a chronic condition, and we next investigated if the NG^5ht3aR^ neurons are required for long-term blood pressure homeostasis. To achieve this, we selectively ablated NG^5ht3aR^ neurons by bilaterally injecting a Cre-dependent AAV expressing Caspase 3 (AAV-DIO-Cas) or a fluorescent Control (AAV-DIO-tdTomato, see Extended Data for details on constructs) in the NG of 5HT3aR^Cre^ rats (Fig. 2G). Using RT-qPCR, we confirmed that Caspase treatment ablated ∼75% of 5HT3aR-expressing neurons in the NG (Fig. 2H). The selective Caspase-dependent ablation of NG^5ht3aR^ neurons produced a sustained time-dependent increase in blood pressure peaking at week 8 compared to the Control group, with no major effects on heart rate (Fig. 2I and Extended Data Fig. 2C). Our findings demonstrate a novel and causal role for a vagal afferent circuit in long-term maintenance of blood pressure homeostasis via the NG^5ht3aR^ neuronal signaling.

### Blunted vagal sensitivity to gut serotonin in hypertension

Because colonic 5-HT was reduced in rodent hypertension following transplant of hypertensive microbiota, we next tested whether vagal sensing of 5-HT may be impaired in hypertension. We first examined 5HT3aR expression in the NG of WKYèWKY and SHRèWKY rats and their respective gut microbiota donors, the WKY and SHR. Using RT-qPCR, we observed diminished 5HT3aR expression in the SHRèWKY as well as their donor SHR along the gut-brain vagal axis. Specifically, expression was reduced in the NG (Fig. 3A) as well as in the peripheral and central sites of vagal terminals in the proximal colon and NTS, respectively (Extended Data Fig. 3A-B). To determine if reduced 5HT3aR expression translated to reduced vagal 5-HT sensing in hypertensive rodents, we investigated the responses of vagal sensory neurons to administration of a specific 5HT3aR agonist L-phenylbiguanide. Using *in vivo* calcium imaging and telemetry allowed us to simultaneously monitor real-time activity of multiple vagal sensory neurons and correlate to blood pressure responses (Fig. 3B) in anaesthetized WKY and SHR. To achieve this, we performed a bilateral microinjection of AAV-hSyn-GCAMP6s into the NG of WKY and SHR to express the calcium indicator GCAMP6 broadly in vagal sensory neurons to serve as a proxy for neuronal activity. We then used two photon microscopy to quantify fluorescence intensity of NG neurons responsive to the infusion of L-phenylbiguanide (Fig. 3B) in anesthetized WKY and SHR with preserved vagal peripheral connections. Vagal sensory neurons in normotensive WKY rats displayed robust dose-dependent responses to L-phenylbiguanide (Fig. 3C, Extended Data Fig. 3C and 3E). This was evident by activation of a significant percentage of responsive NG neurons (Fig. 3C, Extended Data Fig. 3C and E) and a higher GCaMP6s signal amplitude in the WKY compared to the SHR (Fig. 3D bottom panel) that was time-locked to a differential decrease in blood pressures between the strains (Fig. 3D top panel). The neuronal responses were dose-dependent, with a higher number of NG neurons activated at higher L-phenylbiguanide concentrations (Fig. 3C, Extended Data Fig. 3C-E). Notably, the activation persisted for several minutes after the infusion ended (Fig. 3C). In contrast, vagal sensory neurons from SHR exhibited significantly weaker responses to L-phenylbiguanide. A smaller proportion of neurons were activated (Fig. 3C, Extended data Fig. 3C-E), and the overall amplitudes of both the neuronal and blood pressure responses were diminished compared to the WKY rats (Fig. 3D and Extended data Fig. 3C-E).

**Fig. 3:**
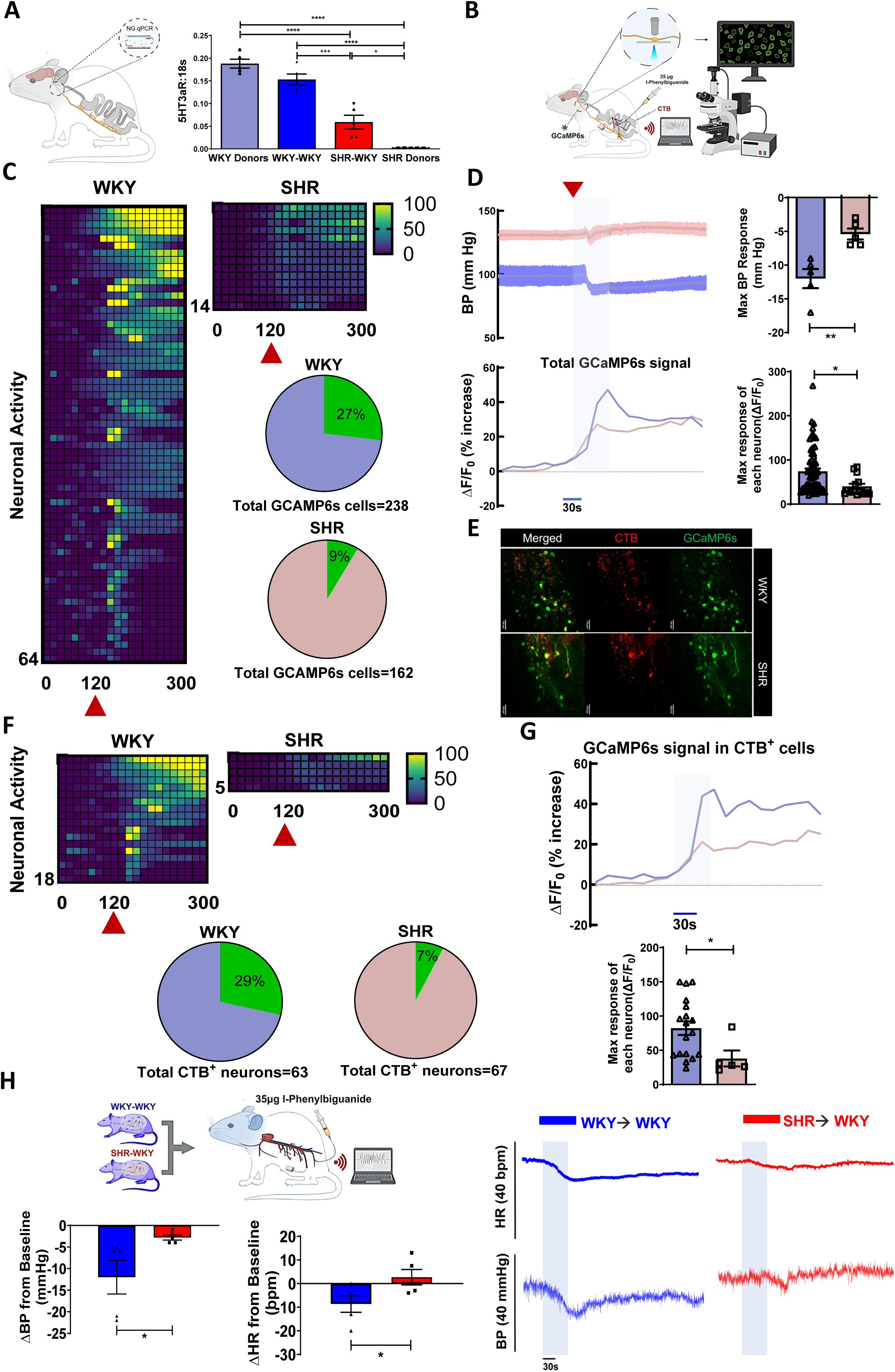
Reduced serotonin signaling by intestinal vagal afferents in hypertensive rodents with gut dysbiosis. **A**, cartoon schematics (created in Biorended) highlights the quantitative real time PCR (qRT-PCR) analysis of relative expression levels of serotonergic 5HT3a receptors (5HT3aR) normalized to 18S housekeeping gene in the nodose ganglia of all donor and recipient groups. A significant reduction in relative expression levels of 5HT3aRs was observed in the nodose ganglia of SHR→WKY group (red) compared to WKY→WKY controls (blue) (by ∼70%), reflecting a similar trend observed in WKY and SHR as their respective gut bacteria donors. Values are means ± SEM; N=5/group; ANOVA with Mann-Whitney post-hoc, *P<0.05, ***P<0.001, ****P<0.0001. **B,** a cartoon schematic (created in Biorender) illustrating *in vivo* recordings of calcium responses in vagal afferents co-expressing GCaMP6s and the red fluorescent cholera toxin B (CTB) expressed retrogradely from the proximal colon, in response to i.v. infusion of 5HT3a receptor (5HT3aR) agonist L-phenylbiguanide. Blood pressure responses were measured simultaneously with neuronal responses in all rats. **C,** the heat maps depict z-score-normalized fluorescence traces from all GCaMP6s-expressing vagal neurons in the nodose ganglia of normotensive Wistar-Kyoto (WKY) rats (left panel) and spontaneously hypertensive rats (SHR, right panel), responding to i.v. injection of the serotonin receptor agonist. Each row represents the activity of a single cell over 5 mins. Stimulus was given at 120 seconds (marked by red triangle). In N=5 per group, 27% of the total of 238 GCaMP6s-expressing neurons responded to serotonin agonist in the WKY rats, with only 9% of total of 162 GCaMP6s-expressing neurons were responsive in the SHR, as depicted by pie charts. The responses of each neuron are represented in the heat maps. **D**, in top left panel, average traces of blood pressure (BP, mmHg) responses to infusion of serotonin receptor agonist in anaesthetized WKY (blue) and SHR (red) (N=6/group). A significantly diminished BP response was observed in the SHR compared to the WKY rats (top right panel). Absolute values are normalized to values observed at baseline for each rat and presented as change (Δ) from baseline. The BP responses were time-locked to the calcium responses of total GCaMP6s-expressing vagal neurons (bottom panel). Neuronal responses are normalized to values observed at baseline (F_0_) for each rat and presented as change (Δ) from baseline (%increase in activity). Average peak amplitudes of vagal responses were also reduced in the SHR (N=5/group). Values are means ± SEM; Student’s Ttest, *P<0.05, **P<0.01. **E**, an image depicting colon-projecting vagal afferents retrogradely labeled by CTB (in red) and total GCaMP6s-expressing nodose ganglia neurons (in green) showing a similar population of neurons in WKY and SHR (also quantified in the pie charts in C and F). Scale bar = 100µm. **F**, the heat maps depict z-score-normalized fluorescence traces from colon-projecting GCaMP6s vagal neurons in the nodose ganglia of normotensive Wistar-Kyoto (WKY) rats (left panel) and spontaneously hypertensive rats (SHR, right panel), responding to i.v. infusion of serotonin receptor agonist. Each row represents the activity of a single cell over 5 mins. Stimulus was given at 120 seconds (marked by red triangle). In N=5 per group, a similar population of neurons is double labeled in the WKY and SHR, and 29% of the total of the 63 colon-projecting GCaMP6s-expressing neurons responded to the serotonin receptor agonist in the WKY rats, with only 7% of the 67 colon-projecting GCaMP6s-expressing neurons were responsive in the SHR, as depicted by pie charts. The responses of each neuron are represented by heat maps. **G**, in top panel, average traces of calcium responses of GCaMP6s-expressing vagal neurons projecting to the colon in anaesthetized WKY (blue) and SHR (red) (N=6/group) in response to i.v. infusion of serotonin receptor agonist L-phenylbiguanide. Absolute values are normalized to values observed at baseline (F_0_) for each rat and presented as change (Δ) from baseline (%increase in activity). Average peak amplitudes of colon-projecting vagal afferents were reduced in the SHR (bottom panel, N=5/group). Values are means ± SEM; Student’s Ttest, *P<0.05. **H**, on top left, cartoon schematic (created in BIorender) illustrating BP (mmHg) and HR (bpm) measurements in anaesthetized WKY→WKY (blue) and SHR→WKY rats (red) in response to i.v. infusion of serotonin receptor agonist L-phanylbiguanide. On the right, representative traces of BP and HR responses in both groups, with shaded bar representing length of infusion. Bottom left panels illustrate average BP and HR peak responses, significantly reduced in the SHR→WKY group (N=5/group). Absolute values are normalized to values observed at baseline for each rat and presented as change (Δ) from baseline. Values are means ± SEM; Student’s Ttest, *P<0.05.

We next investigated if colon-projecting vagal sensory neurons also exhibited reduced responses to 5-HT in hypertension. We first confirmed that a similar subset of colon-projecting NG neurons was labeled in response to injecting a red fluorescent retrograde neuronal tracer cholera toxin B (CTB) in the submucosal wall of the proximal colons of the WKY and SHR (Fig. 3E). When stimulated with L-phenylbiguanide, the colon-specific vagal sensory neurons in WKY rats displayed significantly stronger responses compared to those observed in the SHR (Fig. 3F-G and Extended Data Fig. 3D-F). This was also reflected in a reduced percentage of CTB-labeled neurons that responded to L-phenylbiguanide in the SHR compared to the WKY (Fig. 3F and Extended Data Fig. 3D-F). Because FMT from SHR to WKY reduced colonic 5-HT (Fig. 1C) and the 5HT3aR expression along the vagal gut-brain axis (Fig. 3A and Extended Data Fig. 3A-B), we next assessed if this is linked to blood pressure homeostasis. Indeed, the rapid and sustained reduction in blood pressure and heart rate by L-phenylbiguanide demonstrated in the WKYèWKY was significantly blunted in the SHRèWKY rats (Fig. 3H), mimicking the blood pressure responses observed in the WKY and SHR (Fig. 3D, top panel), and reflective of reduced 5HT3aR expression along the gut-brain vagal axis in hypertension (Fig. 3A and Extended Data Fig. 3A-B). These findings demonstrate a link between pro-hypertensive microbiota and diminished 5-HT vagal sensing in the colon.

### A critical role for colonic vagal serotonin signaling in blood pressure homeostasis and response to stress

Next, we aimed to address the direct and causative role of colonic 5-HT sensing NG neurons in blood pressure homeostasis. First, to confirm that colon-projecting vagal afferents express 5HT3aRs, we injected a retrograde control virus expressing Cre-dependent tdTomato into the wall of the proximal colons of 5HT3a^Cre^ rats (Fig. 4A). We found that the 5HT3aR-expressing vagal sensory neurons extensively innervated the proximal colon and the NTS (Extended Data Fig. 4A). Next, we assessed whether this subpopulation of colonic NG^5HT3aR^ neurons played a role in regulation of blood pressure. To stimulate colonic NG^5HT3aR^ neurons, a Cre-dependent retrograde virus expressing the light sensitive ChR2 (AAVrg-DIO-ChR2, see Extended Data for details on viral vectors) was injected in the wall of the proximal colon, with control animals receiving a Control virus with tdTomato and no ChR2 (Fig. 4A and Extended Data Figure 4A). When we optically and bilaterally stimulated the cell bodies of the colonic NG^5HT3aR^ neurons expressing ChR2, this produced a rapid reduction in blood pressure (Fig. 4B and 4C) at a similar amplitude that was observed before, when NG^5HT3aR^ neurons were stimulated broadly (Fig. 2C and 2E). In contrast, the average amplitudes of heart rate responses to stimulation of the colonic NG^5HT3aR^ neurons (Fig. 4B and 4D) were smaller than those observed following broad stimulation of NG^5HT3aR^ neurons (Fig. 2C and 2F), suggesting that a subset of NG^5HT3aR^ neurons not innervating the colon are involved in regulation of heart rate. Control animals exhibited no changes in response to light (Fig. 4B-D). These findings reveal a previously unknown role for colonic vagal serotonin signaling in short-term regulation of blood pressure.

**Fig. 4:**
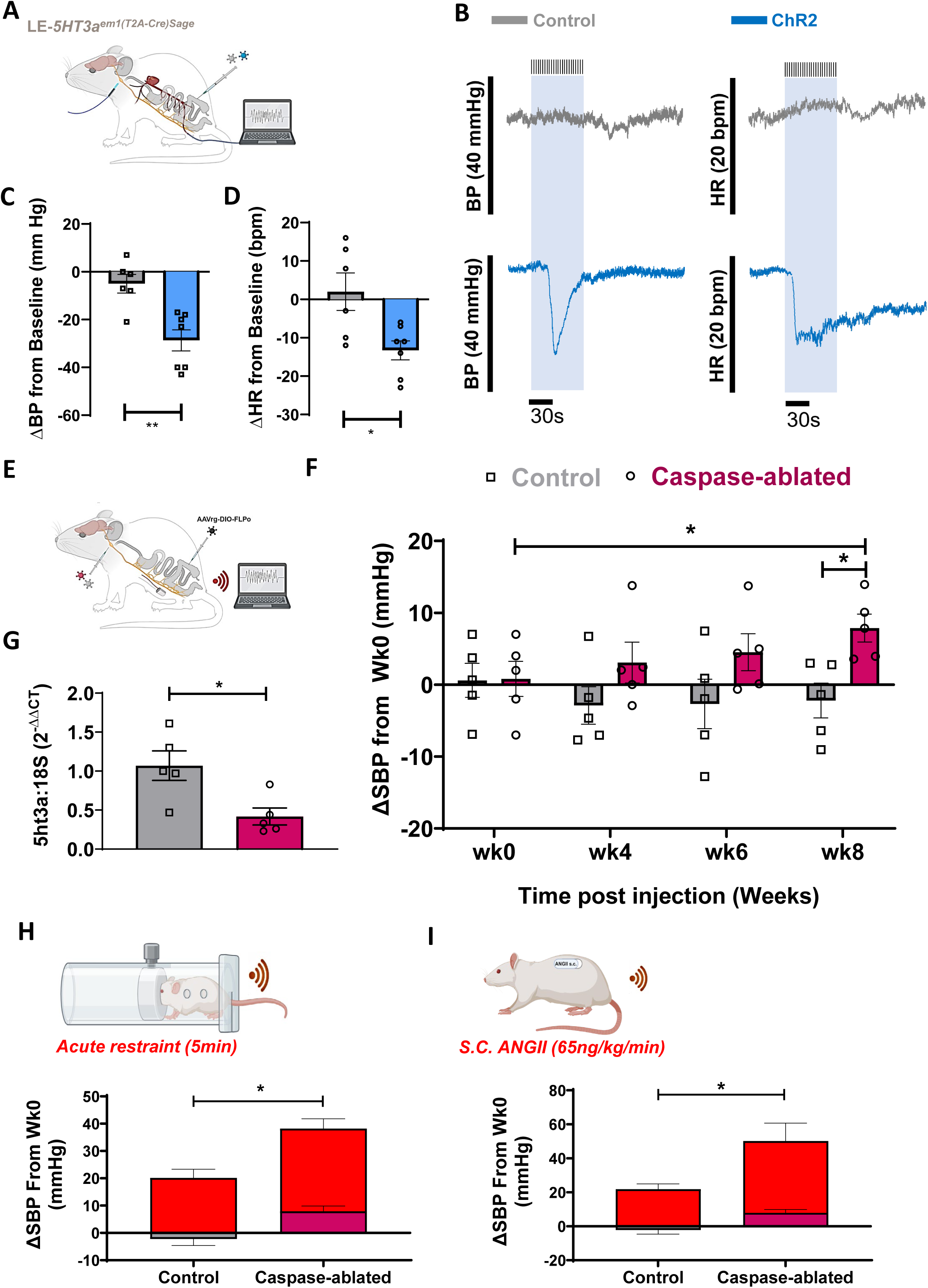
Colonic vagal serotonin signaling mediates blood pressure homeostasis and stress response. **A,** retrograde Cre-dependent viral vectors (AAVrg) expressing either excitatory Channelrhodopsin 2 (ChR2) or Control tdTomato fluorescent protein were microinjected in the proximal colons of LE-5HT3a^em1(T2A-Cre)Sage^ (5HT3aR^Cre^) rats. Three weeks later, rats were anaesthetized, and femoral artery was cannulated for measurement of direct blood pressure (BP, mmHg) and heart rate (HR, beats per minute, bpm) before, during and after application of the blue excitatory light to the nodose ganglia for 1 minute. **B**, representative traces of BP and HR before, during and after the optic stimulation of the nodose ganglia. Optic stimulation of the nodose ganglia expressing ChR2 produced a time-locked decrease in BP and HR (in blue), while optic stimulation of the Control nodose ganglia (in gray) did not produce any significant responses in 5HT3aR^Cre^ rats. Light blue box illustrates timing of optic stimulation. **C**, average BP (mmHg) responses to bilateral optic stimulation of the nodose ganglia expressing ChR2 (blue bar) and Control (gray bar) (N=6-7 rats per group). Absolute BP values following stimulation of each nodose ganglia separately are normalized to values observed at baseline, averaged for each rat, and presented as change (Δ) from baseline. Values are means ± SEM; Student’s Ttest, **P<0.01. **D**, average HR (bpm) responses to bilateral optic stimulation of the nodose ganglia expressing ChR2 (blue bar) and Control (gray bar) (N=6-7 rats per group). Absolute HR values following stimulation of each nodose ganglia separately are normalized to values observed at baseline, averaged for each rat, and presented as change (Δ) from baseline. Values are means ± SEM; Student’s Ttest, *P<0.05. **E,** radiotelemetry transmitters were implanted in femoral artery of 5HT3aR^Cre^ rats for continuous measurement of BP and HR in conscious unrestrained rats. At the same time, a retrograde Cre-dependent vector (AAVrg) expressing FLPo recombinase was injected in colons of all rats. Following recovery and baseline BP recordings (wk0), FLPo-dependent vectors expressing either Caspase 3, to selectively ablate 5HT3aR-expressing vagal afferents that project to the colon only, or Control YFP protein were bilaterally microinjected in the nodose ganglia of separate 5HT3aR^Cre^ rats. Schematics created in Biorender. **F**, radiotelemetric measurements of continuous systolic BP (SBP, mmHg) were performed once a week over 24hrs in conscious unrestrained 5HT3aR^Cre^ rats at baseline (i.e., before nodose ganglia viral infections, noted as wk0) and then at weeks 4-8. A significant increase in SBP (by∼9mmHg) was observed in the Caspase-ablated rats (magenta) compared to Control group (gray) and compared to its baseline (wk0) at wk8 post injection (N=5/group). Absolute values are normalized to values observed at baseline for each rat and presented as change (Δ) from baseline. Values are means ± SEM; ANOVA with Mann-Whitney post-hoc, *P<0.05. **G,** at endpoint, virally mediated ablation of 5HT3aR-expressing colonic vagal neurons was confirmed by reduced relative expression levels of nodose ganglia 5HT3aRs in Caspase-ablated rats (magenta) as measured by quantitative real time PCR (qRT-PCR). N=5/group. Values are means ± SEM; Student’s Ttest, *P<0.05. **H**, BP responses to 5 minutes of acute restraint were measured by radiotelemetry in both sets of rats at wk8, post-injection. 5HT3a^Cre^ rats with Caspase-ablated 5HT3aR-expressing colonic vagal neurons (magenta) presented with significantly higher SBP (mmHg) responses to the acute restraint compared to the Control group (gray). Absolute values are normalized to values observed at w0 for each rat and presented as change (Δ) from wk0, with red bars representing additive effects of restraint at wk8. Values are means ± SEM; ANOVA with Mann-Whitney post-hoc, *P<0.05. **I**, BP responses to a one week of subcutaneous angiotensin II (s.c. ANGII) infusion were measured by radiotelemetry in both groups starting from wk8 post-injection. 5HT3a^Cre^ rats with Caspase-ablated 5HT3aR-expressing colonic vagal neurons (magenta) presented with significantly higher SBP (mmHg) responses to one week of s.c. ANGII infusion compared to the Control group (gray). Absolute values are normalized to values observed at wk8 for each rat and presented as change (Δ) from wk8, with red bars representing additive effects of ANGII. Values are means ± SEM; ANOVA with Mann-Whitney post-hoc, *P<0.05.

To investigate the direct role of colonic NG^5HT3aR^ neurons in long term blood pressure homeostasis, we utilized an intersectional genetic strategy in which the 5HT3aR^Cre^ rats received colonic injections of Cre-dependent retrograde virus to selectively express FlpO recombinase (AAVrg-DIO-FlpO; Fig. 4E, see Extended Data for details on viral constructs) in colonic NG^5HT3aR^ neurons. Following this, we bilaterally injected a new FlpO-dependent viral construct that expresses Caspase (AAV-fDIO-Cas, see Extended Data for details on viral constructs) into the NG (Fig. 4E) to facilitate selective Caspase-mediated ablation of colonic NG^5HT3aR^ neurons only. In control animals, same approach was used to express a control fluorescent protein (YFP) in colonic NG^5HT3aR^ neurons (Fig. 4E). This resulted in viral expression in the colonic NG^5HT3aR^ neurons (Extended data Fig. 4B) and a ∼55% reduction in 5HT3aRs in the NG of 5HT3aR^Cre^ rats injected with the new Caspase construct (Fig 4G). Caspase-mediated ablation of colonic NG^5HT3aR^ signaling significantly increased blood pressure with time, peaking at week 8 post transfection, with no major differences in heart rate between the groups (Fig. 4F and Extended data Fig. 4C). This was independent of gut bacterial composition and abundances (Extended Data Fig. 4D) and demonstrates a crucial role for intact colonic NG^5HT3aR^ signaling in maintaining the blood pressure homeostasis.

Stress is intricately linked with hypertension and gut dysbiosis [29] and the microbiota-vagal axis mediates stress responses [30]. Thus, we tested whether ablation of colonic NG^5HT3aR^ neurons would enhance stress-dependent elevation in blood pressure. We first measured the effects of acute restraint on blood pressure in Control rats and those with Caspase-ablated colonic NG^5HT3aR^ neurons and demonstrate that rats with selective ablation of colonic NG^5HT3aR^ neurons exhibited enhanced stress-induced blood pressure responses compared to Controls (Fig. 4H). We next tested if administration of angiotensin II (ANGII), a key pro-hypertensive hormone intricately linked with stress and hypertension [31], may promote hypertension in rats with selective ablation of colonic NG^5HT3aR^ neurons. We observed that ANGII infused continuously over one week further enhanced hypertension in rats with selective ablation of colonic NG^5HT3aR^ neurons (Fig. 4I). These data demonstrate that intact intestinal serotonergic vagal signaling is necessary for blood pressure homeostasis and that it can mediate stress responses in rats.

### Recovery of vagal serotonergic signaling in the colon can alleviate dysbiosis-induced hypertension

Because hypertensive microbiota can diminish 5HT3aR signaling, we assessed whether reinstating the expression of 5HT3aRs in colonic NG^5HT3aR^ neurons could alleviate hypertension induced by gut dysbiosis. In the hypertensive offspring of SHR dams and 5HT3aR^Cre^ males (SHR/5HT3aR^Cre^ rats) that present with gut dysbiosis (Extended data Fig. 5A-B), we selectively overexpressed 5HT3aRs in colonic NG^5HT3aR^ neurons using a similar intersectional approach as before (Fig. 4E). For this, a Cre-dependent retrograde virus expressing FlpO recombinase was injected into the colonic wall of SHR/5HT3aR^Cre^ rats (Fig. 5A). Subsequently, a second FLpO-dependent virus (AAV-fDIO-5HT3aR) was injected bilaterally into the NG to selectively overexpress 5HT3aRs in colonic NG^5HT3aR^ neurons (Fig. 5A). A control virus expressing a fluorescent protein (AAV-fDIO-YFP) was injected bilaterally into NG in the control group (Fig. 5A). This targeted approach resulted in fluorescent viral labeling in a subset of colonic NG^5HT3aR^ neurons in both groups (Fig. 5B) and elevated 5HT3aR expression by ∼30% in the NG of SHR/5HT3aR^Cre^ rats injected with a virus expressing 5HT3aRs, as confirmed by RT-qPCR (Fig. 5C).

**Fig. 5.**
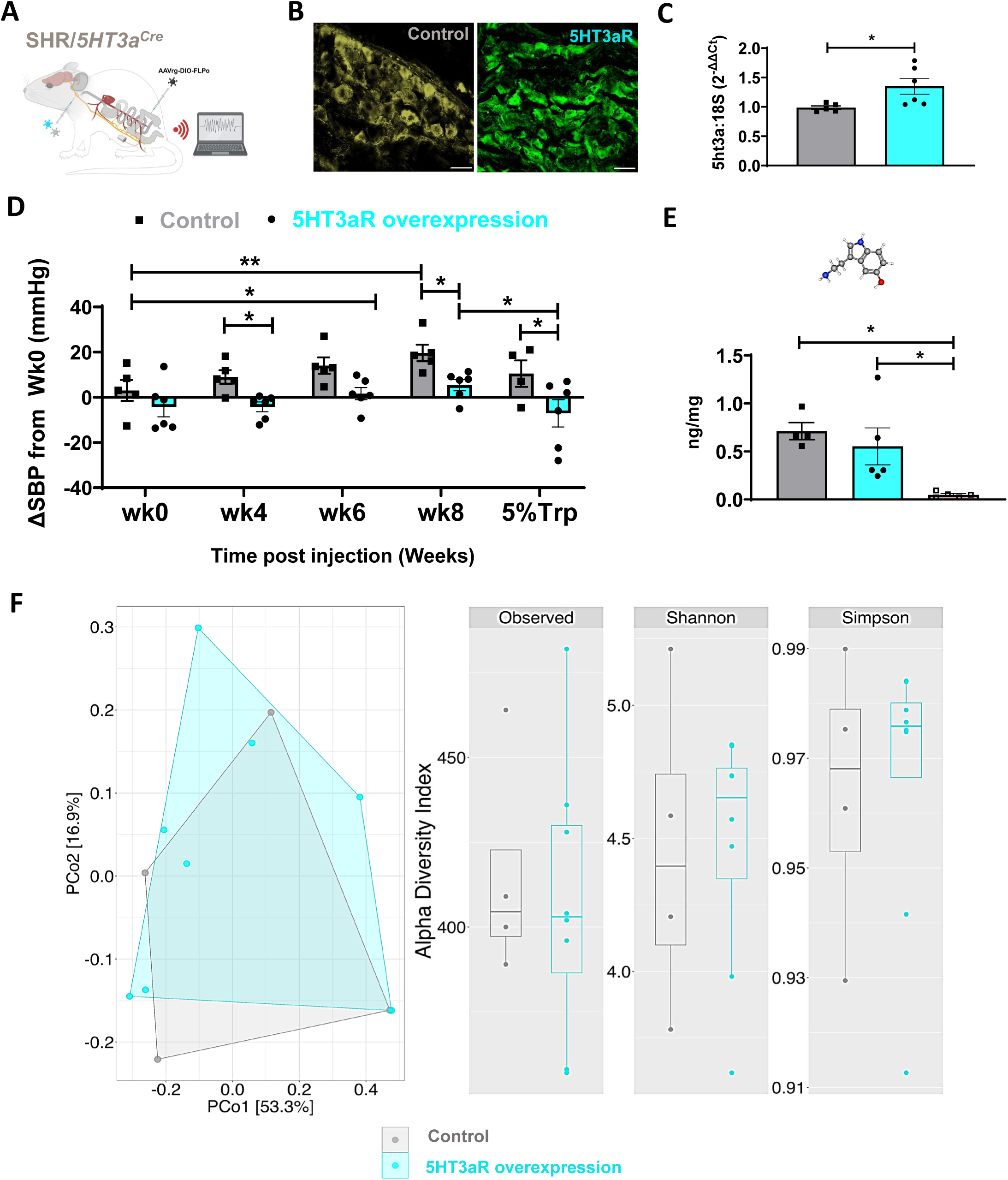
Restoring gut vagal serotonin signaling alleviates microbiota-induced hypertension. **A,** radiotelemetry transmitters were implanted in femoral artery of hypertensive male and female SHR/5HT3aR^Cre^ rats for continuous measurement of blood pressure (BP, mmHg) and heart rate (HR, beats per minute, bpm) in conscious unrestrained rats. At the same time, a retrograde Cre-dependent vector (AAVrg) expressing FLPo recombinase was injected in colons of all rats. Following recovery and baseline BP recordings (wk0), FLPo-dependent vectors (AAV) expressing either the 5HT3a receptors (5HT3aRs) or control YFP fluorescent protein were bilaterally microinjected in the nodose ganglia in separate rats. Schematics created in Biorender. **B**, endpoint confirmation of viral expression of control YFP (in yellow) and 5HT3aR (in green) in the nodose ganglia of SHR/5HT3aR^Cre^ rats. Scale bar=100µm. **C,** at endpoint, virally mediated overexpression of 5HT3aRs was confirmed by quantitative real time PCR (RT-qPCR) in the nodose ganglia. N=6/group. Values are means ± SEM; Student’s Ttest, *P<0.05. **D**, radiotelemetric measurements of continuous systolic BP (SBP, mmHg) were performed once a week over 24hrs in conscious unrestrained SHR/5HT3aR^Cre^ rats at baseline (i.e., before nodose ganglia viral injections, noted as wk0) and then at weeks 4-8. Expression of 5HT3aRs (cyan) in colonic vagal neurons significantly alleviated time-dependent increase in SBP compared to the Control group (gray). This effect was potentiated by dietary supplementation with 5% tryptophan (Trp) administered for one week *ad libitum*, but only in rats with 5HT3aR overexpression (in cyan). Absolute values are normalized to values observed at baseline for each rat and presented as change (Δ) from baseline. N=4-6/group. Values are means ± SEM; ANOVA with Mann-Whitney post-hoc, *P<0.05, **P<0.01. **E**, one weeklong supplementation with 5% Trp in diet elevated colonic serotonin in both the Control (in gray) and 5HT3aR overexpression (in cyan) groups when compared to the SHR (light red, open squares). N=5/group. Values are means ± SEM; ANOVA with Mann-Whitney post-hoc, *P<0.05. **F**, endpoint 16S bacterial sequencing analyses show an overlap in composition and abundances of the gut bacteria between the two groups, illustrated by the principal coordinate analysis (PCoA) showing beta diversity with Bray-Curtis Dissimilarity (left) and Alpha diversity plots with the common indices (right) (N=4-6/group).

Remarkably, the targeted increase of 5HT3aRs on colonic vagal afferents mitigated the development of hypertension (Fig. 5D and Extended Data Fig. 5C). This was further potentiated by supplementation with dietary Trp (Fig. 5D), which elevated colonic 5-HT (Fig. 5E) independently of gut dysbiosis (Fig. 5F). This demonstrated that colonic 5HT3aR-dependent vagal signaling is a crucial link between the gut microbiota and blood pressure homeostasis and that recovery of this signaling can mitigate gut dysbiosis-associated hypertension.

## Discussion

Our study characterizes a previously unknown role of intestinal vagal sensory neurons in orchestrating the communication from the gut microbiota to maintain the blood pressure homeostasis. Our investigation delineates a distinct population of colonic vagal sensory neurons responsive to 5-HT whose activity is reduced in hypertension, subject to influence of the gut microbiota. Recent studies have demonstrated a role for gut dysbiosis in development of high blood pressure [7, 13] but the direct causative mechanisms were unclear. The vagus nerve is a master regulator of an array of homeostatic functions including by mediating the communication between the gut bacteria and its host [32]. Our novel findings extend the current scientific knowledge to include a critical role for the intestinal vagus in blood pressure homeostasis via the serotonin signaling. Our findings complement the recent reports of intricate interplay between the gut microbiota and intestinal 5-HT [33–35] and those linking deregulation in Trp metabolism with human and rodent hypertension [15–17] by demonstrating a new neural link between the gut bacteria and host intestinal serotonin and hypertension. As such, these findings may have significant implications for the development of future anti-hypertensive therapeutics.

We utilized an intersectional genetic strategy and new viral tools to underscore the pivotal role of 5-HT-sensing intestinal vagal neurons in regulation of blood pressure. However, precise identity of central networks involved in intestinal regulation of blood pressure is unknown. Notably, prior research has implicated select NTS neurons in governing the 5-HT vagal signaling originating from the gut with evidence of influence over several homeostatic functions including feeding and psychological stress [36–41]. We posit that a substantial degree of signaling convergence exists among several processes within the NTS and beyond, which will be investigated in future studies. The overlap between feeding and blood pressure may be particularly relevant as intestinal 5-HT, released in response to ingestion of nutrients to promote digestion, secretion, and, motility can also reduce the activity of the sympathetic nervous system and promote intestinal vasodilation [42] and postprandial hyperemia [43]. Indeed, GI dysfunction and diminished GI blood flow has been reported in the SHR [8], while constipation is reported in some hypertensive patients [44, 45] further linking the gut dysbiosis with GI pathology in hypertension.

We observed a significant potentiation of pro-hypertensive responses during stress following disruption of intestinal 5-HT vagal signaling. This is in alignment with emerging role of the gut microbiota-vagal axis and Trp metabolism in conditions of stress [46] and with findings of exaggerated blood pressure responses to pro-hypertensive FMT under restraint [12, 13]. This highlights the intestinal microbiota-gut-vagal serotonin signaling as pivotal in blood pressure homeostasis as well as in blood pressure responses to daily cardiovascular stressors.

Reduced expression of colonic Tph1 following the pro-hypertensive FMT correlated with reduced colonic 5-HT in same rats. Moreover, no change in either the colonic or bacterial Trp or bacterial 5-HT demonstrates that reduced colonic 5-HT is a function of direct gut bacterial effects. Gut bacteria can stimulate the intestinal 5-HT via butyrate-induced modification of Tph1 expression [47, 48]. Thus, reduced fecal butyrate following transplant of the pro-hypertensive bacteria may be driving intestinal 5-HT depletion in our model but this remains to be directly tested. Gut microbiota composition is highly influenced by external factors such as the diet and antibiotic treatment as well as internal factors that govern the host intestinal environment. Diet especially is a likely culprit in today’s climate of increased consumption of Westernized diets, and dietary interventions in combination with select butyrate-stimulating pre- and probiotics [49] may present a viable therapeutic intervention in combination with currently available antihypertensive medication to alleviate the burden of treatment resistant hypertension.

Resistant hypertension, characterized by unresponsiveness to three or more antihypertensive medications [50], is present in up to 20% of hypertensive patients [4–6, 51]. The currently available drugs are designed to target cardiovascular, neural, and renal mechanisms, and as such may be lacking influence on hypertension driven from the gut and its microbiota. Moreover, the presence of gut dysbiosis has been associated with unresponsiveness to commonly used antihypertensive medications in individuals with resistant hypertension [52] demonstrating the need for medicine that will incorporate the effects on and by the gut microbiota. Our findings deepen the understanding of complex dynamics governing host-microbiota interactions and signal a paradigm shift in antihypertensive therapeutic inquiry, paving the way for innovative treatments targeting 5HT3a receptor agonism in tandem with dietary, pre-, and pro-biotic interventions. Such endeavors offer promising avenues for the development of precision therapeutics aimed at reinstating physiological equilibrium and alleviating the burden of resistant hypertension associated with gut dysbiosis.

### Online content

Additional source data, extended data, supplementary information, and statements of data and code are available at PRJNA1140871 http://www.ncbi.nlm.nih.gov/bioproject/1140871.

### Grants

Research reported in this publication was supported by National Heart, Lung, and Blood Institute of the National Institutes of Health under award number R01HL152162-01A

## Methods

Details on antibodies, viral vectors, and Trp diet are found in the Extended Data.

### Experimental rat models

All procedures were approved by the Institutional Animal Care and Use Committee at the University of Toledo and University of Florida. Studies were performed in male and female rats aged 2-5 months. Rats were group housed until telemetry surgery in a specific pathogen-free facility (22°C, 65% humidity and 12h light:dark cycle) with free access to regular chow and systems water. Strains used were as follows: male WKY (normotensive, Charles River), male and female SHR (hypertensive, Charles River), and male and female HsdSage:LE-*5ht3a*^em1(T2-Cre)Sage^ (normotensive, 5HT3aR^Cre^, Inotiv). To generate SHR/5ht3aR^Cre^, female SHR was cross bread with male 5HT3aR^Cre^. FMT experiments were performed on male WKY (by oral gavage, as per [11]) and on male and female SHR/5ht3aR^Cre^ offspring by biweekly transfer of feces in bedding from adult sex matched SHR donors [53]) starting at weaning until 3 months of age. Success of FMT was confirmed by 16S rRNA sequencing and analysis [54].

### Metabolomic analyses in colons and feces

Samples were homogenized at 50 mg/mL in 5 mM Ammonium acetate. Protein concentration of colon samples was normalized to 1100 ug/mL, and fecal samples were normalized per g of wet feces. Global metabolomics profiling was performed on a Thermo Q-Exactive Orbitrap mass spectrometer with Dionex UHPLC. Samples were analyzed in positive and negative heated electrospray ionization with a mass resolution of 35,000 at m/z 200 as separate injections. Separation was achieved on an ACE 18-pfp 100 x 2.1 mm, 2 µm column with mobile phase A as 0.1% formic acid in water and mobile phase B as acetonitrile. The column flow rate was 350 µL/min at 25°C. 4 µL was injected for negative and 2 µL for positive ions. MZmine (freeware) was used to identify features, deisotope, and align features. All adducts and complexes were identified and removed from the data set. The mass and retention time data was searched against our internal metabolite library [55, 56]. Inner-quartile range filtering was implemented in the R package MetaboAnalystR (https://github.com/xia-lab/MetaboAnalystR). Missing data were imputed by k-nearest neighbor imputation in the R package MetaboAnalystR. Peak intensities were normalized sample-wise using sum normalization followed by log (base 10) transformation and Pareto scaling in R package MetaboAnalystR. Multivariate and univariate statistical analysis assessed clustering of samples based on metabolomic profiles to identify metabolites with changes in abundance between groups or biological conditions. The list of compounds detected in positive and negative ionization modes was combined and reduced to a single representative compound per likely metabolite based on p-value (lowest p-value compound retained). For targeted tryptophan metabolomics, the target analytes, L-tryptophan and 5-HT were obtained from Sigma (St Louis, MO), L-kynurenine, kynurenic acid and xanthurenic acid were obtained from MP biomedicals (Santa Ana, CA), and anthranilic acid was obtained from Fluka (Charlotte, NC). L-tryptophan 13C11 and L-kynurenine sulfate:H2O (ring D4) were obtained from Cambridge Isotopes (Tewksbury, MA), serotonin-D4, kynurenic acid-D5 and xanthurenic acid-D4 were obtained from Santa Cruz (Dallas, TX), and anthranilic acid-13C6 and xanthurenic acid-D4 were obtained from ISOTEC (St Louis, MA). All other reagents used were of LC-MS purity and obtained from ThermoScientific. Bovine serum albumin (BSA) (Sigma) was prepared to a concentration of 70 g/L in LC-MS grade water. A stock calibration mix was prepared in BSA with Trp at 80,000 ng/mL, 5-HT and kynurenine at 8,000 ng/mL, kynurenic acid, xanthurenic acid and anthranilic acid at 4,000 ng/mL. A 9-point calibration curve was created with serial dilution covering 100 – 25,600 ng/mL for Trp, 10 – 2,560 ng/mL for 5-HT and kynurenine, and 5 – 1,280 for kynurenic acid, xanthurenic acid and anthranilic acid. The internal standard solution was prepared with Trp- 13C11, 5-HT-D4 and kynurenine-D4 at 5 µg/mL, kynurenic acid-D5 at 0.5 µg/mL and anthranilic acid 13C6 and xanthurenic acid-D4 at 0.8 µg/mL in 90:10 water:methanol. Sample analysis was conducted on a ThermoScientific Q-Exactive Orbitrap with Dionex 3000 UHPLC. Briefly, samples were analyzed in positive heated electrospray ionization with the following source parameters: spray voltage = 3500 V, aux gas = 10, sheath gas = 50, capillary temperature = 325 °C, spray gas = 1, and S-lens RF level = 30. Separation was achieved on an ACE 18-PFP 100 x 2.1 mm, 2 µm column (Mac-Mod Analytical, Chadds Ford, PA) using a gradient with mobile phase A as 0.1% formic acid in water and mobile phase B as acetonitrile with a column temperature of 25°C. Gradient elution was ramped from 0% B to 80% B over 13.0 min at 350 µL/min and increased to 600 µL/min from 16.80 and 17.50 min for re-equilibration. The runtime was 20.50 min and full scan at 35,000 mass resolution was acquired from 2 µL injection in positive ion mode. Calibration was conducted by extraction of the exact mass for each target analyte and reference to the respective internal standard.

### Serotonin (5-HT) assay in the proximal colon

5-HT ELISA was performed at endpoint in proximal colons of adult male WKY and SHR, and male and female SHR/5HT3aR^Cre^ rats following Trp diet administration. Rat proximal colons were dissected at endpoint and flash frozen. Homogenized tissues (300 µl of RIPA buffer Abcam, Cat #:ab156034, with protease and phosphatase inhibitor cocktail, Abcam, Cat #:ab 201120) were used to measure 5-HT levels in technical triplicates using a competitive enzyme-linked immunosorbent assay (ELISA) following the manufacturer’s instructions (Rocky Mountain Diagnostics, CO, USA, Cat # BA E-5900R).

### Bacterial DNA extraction, 16S rRNA sequencing and data analysis

Fecal and/or cecal samples were collected for 16S rRNA gene sequencing as per [54, 57, 58]. Briefly, bacterial DNA extraction was performed using QIAam®PowerFecal®DNA kit (QIAGEN) and bacterial 16S rRNA gene sequencing was performed on an Illumina MiSeq (Illumina, San Diego,CA) platform. Demultiplexed reads were analyzed using QIIME 2 [v.2023.2] [54, 58, 59]. DADA2 plugin was used to merge paired-end FastQ files, denoising, chimera removal and construction of amplicon sequence variants (ASV) [60]. Taxonomy was analyzed using the Silva v.132 databases at 99% identity pre-trained for the V3-V4 region [61]. Rarefaction was performed at a sampling depth of the sample with the minimum amount of reads per sample. Alpha-diversity was estimated using 3 metrics: Shannon, Observed features and Simpson. Beta-diversity was performed using the Bray Curtis dissimilarity index and visualized using principal coordinate analysis (PCoA). Permutational Multivariate Analysis of Variance (PERMANOVA, adonis, R, 999 permutations) was used to test whether differences between groups [52, 54, 58, 59, 61].

### General surgical procedures

Radiotelemetry surgeries were performed as per [54, 62]. Briefly, rats were anaesthetized with isoflurane (induction at 4%; maintenance at 2% in 100% medical grade O_2_). Radiotelemetry transmitters (HD-S10, DSI) were implanted in the abdominal or femoral arteries for continuous collection of blood pressure and heart rate in conscious unrestrained rats. Following recovery, blood pressure and heart rate measurements were taken once weekly over 24hrs, averaged and presented as change (Δ) from baseline (i.e., week 0, Wk0) within experimental groups.

Viral vectors were delivered in the wall of the proximal colon and bilaterally in the NG as previously described [28, 63]. For NG injections, in anaesthetized rats, a midline incision in the skin of the neck was made and the muscle retracted to visualize the NG. Glass micropipette (WPI Single-Barrel Standard Borosilicate Glass with Filament OD1.5mm, ID0.84mm; 40µm tip beveled at 30°) attached to a picopump (WPI) was used to inject AAV into NG bilaterally (1µl/NG). For GI injections, a glass micropipette (WPI Single-Barrel Standard Borosilicate Glass with Filament OD1.5mm, ID0.84mm; 40µm tip beveled at 45°) attached to a picopump was used to deliver a series of injections into the wall of the proximal colon at a submucosal level (2 µl/rat).

In a group of rats, ANGII (Bachem Catalogue#H-1705, 65ng/kg/min) was delivered continuously via a mini osmotic pump (Alzet 2004) as before [62].

### *In vivo* calcium imaging in the NG of WKY and SHR

*In vivo* calcium imaging was performed in the NG of anaesthetized rats as previously described with some modifications [28]. Male adult WKY and SHR were first injected with choleratoxin B (CTB, Alexa Fluor 594, Thermo Fisher) in the wall of the proximal colon, and with AAV-GCaMP6s bilaterally in the NG as described above. Three weeks later, rats were anaesthetized with isoflurane (induction at 4%; maintenance at 2% in 100% medical grade O_2_). Under anesthesia, rats were implanted with blood pressure catheters in the femoral artery, and with a separate femoral vein catheter (PE, Braintree Scientific R-FC M) that was used for delivery of increasing doses of L-phenylbiguanide (Sigma Aldrich Catalogue#164216, 0-35µg in 150µl of 0.9% NaCl). Rats were kept on a heating pad to maintain body temperature throughout the whole procedure. As described above, the NG and the vagus nerve were dissected, and the vagus nerve was cut immediately above the NG. The NG was gently placed on a stable imaging platform consisted of an 8 mm diameter coverslip attached to a metal arm affixed to a magnetic base. Surgical silicone adhesive (Kwik-Sil, WPI) was applied onto the vagus nerve to immobilize it on the coverslip, and the NG immersed in a drop of DMEM (Corning, 10-014-CV) media was covered with a second coverslip. Imaging was performed using a 16x water immersion, upright objective on a 2-Photon microscope (Bruker) fitted with a Galvanometer for acquisition, and a piezo objective combined with a galvo/resonant scanner. Images were obtained at a frame rate of 30 frames/second. Following baseline measurement of NG GCaMP6s activity and blood pressure (2 minutes), a slow i.v. infusion of L-phenylbiguanide (0-35µg in 150µl saline) was performed using a syringe pump (Harvard apparatus PHD 2000, Cat no. 70-2002) for 1 minute while simultaneously measuring the NG GCaMP6s-dependent green fluorescence and blood pressure responses in real time for 3 minutes. GCaMP6s fluorescent changes were outlined in regions of interests (ROIs) with each ROI defining a single cell throughout the imaging session. The pixel intensity in ROIs (average across pixels) were calculated frame by frame (ImageJ) and exported to Excel and GraphPad Prism for further analyses. The baseline signal was defined as the average GCaMP6s fluorescence over a 2 min period prior to the stimulus introduction. Cells were considered responsive if the following criteria were met: 1) the peak GCaMP6s fluorescence was two standard deviations above the baseline mean, and 2) the mean GCaMP6s fluorescence was 20% above the baseline mean for a 5 second window around the peak. NG in which neurons did not present baseline activity were excluded from the study. GCaMP6s responses in CTB-labeled NG neurons were analyzed separately to represent the activity of colon-projecting vagal afferents. Blood pressure responses were monitored in real time and time locked to GCaMP6s neuronal signal for correlation. At endpoint, NG were extracted and imaged with a Bruker 2-Photon microscope as above for confirmation of viral gene expression.

### Pharmacologic stimulation of 5HT3aRs in WKY**è**WKY and SHR**è**WKY rats

Male WKYèWKY and SHRèWKY rats were anaesthetized with isoflurane (induction at 4%; maintenance at 2% in 100% medical grade O_2_). Under anesthesia, rats were implanted with blood pressure catheters in the femoral artery and a separate femoral vein catheter (PE, Braintree Scientific R-FC M) was used for delivery of L-phenylbiguanide (Sigma Aldrich Catalogue#164216, 35µg in 150µl of 0.9% NaCl). Following baseline measurements (2 minutes), a slow i.v. infusion of L-phenylbiguanide was performed using a syringe pump (Harvard apparatus PHD 2000, Cat no. 70-2002) for 1 minute while simultaneously measuring blood pressure and heart rate in real time. Peak responses were presented as change (Δ) from baseline for each rat and averaged within each experimental group.

### Optogenetic stimulation of NG in 5HT3aR^Cre^ rats

Male and female 5HT3aR^Cre^ rats were injected with viral constructs expressing the control tdTomato or ChR2 either in the wall of the proximal colon, for retrograde expression, or directly in the NG as described above. Three weeks later, rats were anaesthetized (induction at 4%; maintenance at 2% in 100% medical grade O_2_) and femoral blood pressure catheter (PE, Braintree Scientific R-FC M) was implanted, and two ECG leads were used for continuous monitoring of blood pressure and heart rate with Spike2 (CED) under anesthesia. NG was exposed bilaterally as before, and intermittent blue light (470nm, 5Hz) was applied directly to each intact NG separately for one minute while simultaneously measuring blood pressure and heart rate. Peak responses to bilateral NG stimulation were averaged for each rat and presented as change (Δ) from baseline. In one group of 5HT3aR^Cre^ rats, blue stimulating light was applied to the NG while simultaneously recording electrical activity of the vagus rostral to the NG, as previously described [64]. At endpoint, NG were extracted and imaged with a Bruker 2-Photon microscope as before for confirmation of viral gene expression.

### Restraint stress in 5HT3aR^Cre^ rats

A group of adult male 5HT3aR^Cre^ rats were bilaterally injected in the NG with FLpO-dependent AAV expressing either Cas or YFP, and in the proximal colon wall with Cre-dependent AAVrg expressing FLpO as described above. Blood pressure and heart rate were measured at baseline and once weekly for 8 weeks as described above. Following this, rats were exposed to acute restraint stress as in [65]. Briefly, a baseline blood pressure recording was made in unrestrained rats for 20 min by telemetry. Following this, all rats were placed into a restrainer (Kent Scientific) for 5 minutes while simultaneously measuring blood pressure. Data was averaged for the duration of the baseline recording and compared to peak blood pressure responses under restraint. Data is presented as change (Δ) from baseline for each rat and averaged within experimental groups. At endpoint, NGs were extracted for RT-qPCR quantification of 5HT3aRs in both groups (see methods below), and for imaging with a Bruker 2-Photon microscope for confirmation of viral gene expression of yellow fluorescent protein (YFP) in Control group.

### Administration of Trp diet in SHR/5HT3aR^Cre^ rats

A group of adult male and female SHR/5HT3aR^Cre^ rats were bilaterally injected into NG with FLpO-dependent AAV expressing either 5HT3aR or control YFP, and into the proximal colon wall with Cre-dependent AAVrg expressing FLpO as described before. Blood pressure and heart rate were measured in all rats at baseline and then once weekly for eight weeks following viral injections as before. Following this, all rats were administered 5% Trp in diet (Research Diets, Catalogue#A20012701, see Extended Data for dietary content) *ad libitum* for one week. Blood pressure and heart rate were measured by telemetry one week following Trp diet administration. Data was presented as change (Δ) from baseline for each rat and averaged within experimental groups. Food and water intake were measured once a week over 48hrs, prior and during the week of administration of Trp diet. At endpoint, NGs from both groups were extracted and imaged with a Bruker 2-Photon microscope as before for confirmation of viral gene expression as before.

### RNA Isolation and RT-qPCR in the proximal colon, NTS and NG

RNA isolation from proximal colon, NTS and NG was performed as previously described [14, 66]. Real time quantitative PCR (RT-qPCR) was used to measure relative expression levels of 5HT3aRs using the following forward and reverse primers: *F-ACC GCC TGT AGC CTT GAC, R- TGC TCT TGT CCG ACC TCA* (IDT). Expression data were normalized for the expression of housekeeping gene 18S (*F-CAT TCG AAC GTC TGC CCT AT, R-GTT TCT CAG GCT CCC TCT CC*) (IDT). Relative expression levels of Tph1 were measured using the following forward and reverse primers: *F- AGC ATA ACC AGC GCC ATG AA, R- GGC ATC ATT GAC GAC ATC GAG* (IDT). RT-qPCR was performed using a CFX96 Touch Real-Time PCR Detection System (Bio-Rad, Hercules, CA) [14, 66]. Relative gene expression was assessed using the relative ΔΔC_t_ or alternatively ΔC_t_ method for comparisons of more than two groups, where C_t_ represented the threshold cycle.

### Immunohistology and imaging

Tissues were post-fixed in 4% PFA (in PBS, pH=7.4) and cryopreserved in 30% sucrose for several days before being embedded in OCT Compound (Fisher Catalogue#23-730-571) and frozen at -80°C. Tissues were sectioned in a cryostat at 35-50µm. To better visualize viral expression of YFP, some tissues were incubated with chicken anti-GFP primary antibody (1:300, Aves labs, GFP-1010) [67, 68] followed by Goat anti-Chicken IgY Alexa Fluor 488 (1:500, Invitrogen, A-11039). For 5-HT staining in the colon, primary rabbit anti-5-HT (1:500, Immunostar, 20080) and anti-rabbit secondary antibody (1:1000, Invitrogen, A-21245) were used. Sections were first washed in Tris-buffered saline (TBS) with 0.05% Tween-20, followed by incubation with 10% normal serum. Tissues were incubated with appropriate primary antibodies overnight at 4°C, then rinsed in TBS/0.05% Tween-20 solution, followed by incubation with secondary antibodies for 40-60 minutes at room temperature. Slides were mounted (Abcam, ab104139) and cover slipped. Fluorescence imaging was performed with the Leica Microsystems TCS SP5 multi- photon laser scanning confocal microscope (10-40x objective lens, 1-5µm z step size). Pseudo-coloring from green to yellow was performed in ImageJ in tissues expressing viral YFP for clarity.

### Statistical analyses

Sample sizes were determined according to standard power analysis (80% power and α = 0.05) based on Cohen’s d standardized effect sizes as in previous reports [54, 62]. Animals were randomly assigned to treatment groups. Whenever feasible, data collection and/or analyses were performed by a blinded experimenter. After testing for normality (Shapiro–Wilk test), intergroup differences were analyzed by unpaired or paired, two-tailed Student’s t-test for single comparison, or by one-way or two-way analysis of variance (ANOVA) with an appropriate multiple-comparison test, as indicated in the figure legends. If nonparametric testing was indicated, intergroup differences were analyzed by Mann–Whitney U-test or Kruskal–Wallis test with Dunn’s correction, as appropriate. The F test for equality of variances was performed and, if indicated, appropriate corrections were made (Welch’s correction for the t-test) using Prism 9 (GraphPad). Data are presented as mean ± SEM.

## Data availability

Any additional data requests are available from the corresponding authors upon request.

## Acknowledgements

The authors like to thank Allen Schroering at the University of Toledo Integrated Core Facilities for technical support with immunohistochemistry.

## Author contributions

A.d.A., E.D. and T.Y. conducted various experiments, performed data analyses, and edited the manuscript. H.S.K. and A.A.P. performed bacterial sequencing analyses and 5-HT assays. N.B. performed animal husbandry, tissue collections, and conducted RT-qPCR experiments and analyses. N.A. performed RT-qPCR experiments and analyses. B.D.A. performed FMT experiments and blood pressure analyses. D.M.B. performed blood pressure measurements during *in vivo* calcium imaging and edited the manuscript. T.J.G. and C.J.M. performed metabolomics experiments and analyses and edited the manuscript. D.S.M. performed tissue cutting, IHC and imaging. A.S. performed blood pressure measurements and edited the manuscript. A.L.K. and H.S.K. performed confocal imaging and edited the manuscript. G.d.L. and J.Z. conducted the experiments, performed the data analyses, provided funding, and wrote and edited the manuscript.

## Competing interests

None.

## Additional information

None.

## Extended Data File

**Table 1.** List of viral vectors.

| **Viral vector** | **Abbreviation** | **Vendor** | **Catalogue Number** |
| --- | --- | --- | --- |
| AAV9-EF1a-fDIO-taCasp3-TEVp-WPRE | AAV-fDIO-Cas | Vector Biolabs | Custom made (see vector map below) |
| AAV9-Ef1a-fDIO EYFP | AAV-fDIO-YFP | Addgene | 55641-AAV9 |
| AAVrg-EF1a-DIO-FLPo-WPRE-hGHpA | AAVrg-DIO-FLPo | Addgene | 87306-AAVrg |
| AAV9-Syn-GCaMP6s-WPRE-SV40 | AAV-GCaMP6s | Addgene | 100843-AAV9 |
| AAV9-EF1a-DIO-tdTomato | AAV-DIO-tdTomato | Addgene | 28306-AAV9 |
| AAV9-Ef1a-fDIO-5htr3a-2A-eGFP | AAV-fDIO-5ht3aR | Vector Biolabs | Custom made (see vector map below) |
| AAVrg-EF1a-DIO-tdTomato | AAVrg-DIO-tdTomato | Addgene | 28306-AAVrg |
| AAVrg-EF1a-DIO-hChR2(H134R)-eYFP-WPRE-HGHpA | AAVrg-DIO-ChR2 | Addgene | 20298-AAVrg |
| AAV5-flex-taCasp3-TEVp | AAV-DIO-Cas | Addgene | 45580-AAV5 |
| AAV9-EF1a-DIO-hChR2(H134R)-eYFP-WPRE-HGHpA | AAV-DIO-ChR2 | Addgene | 20298-AAV9 |

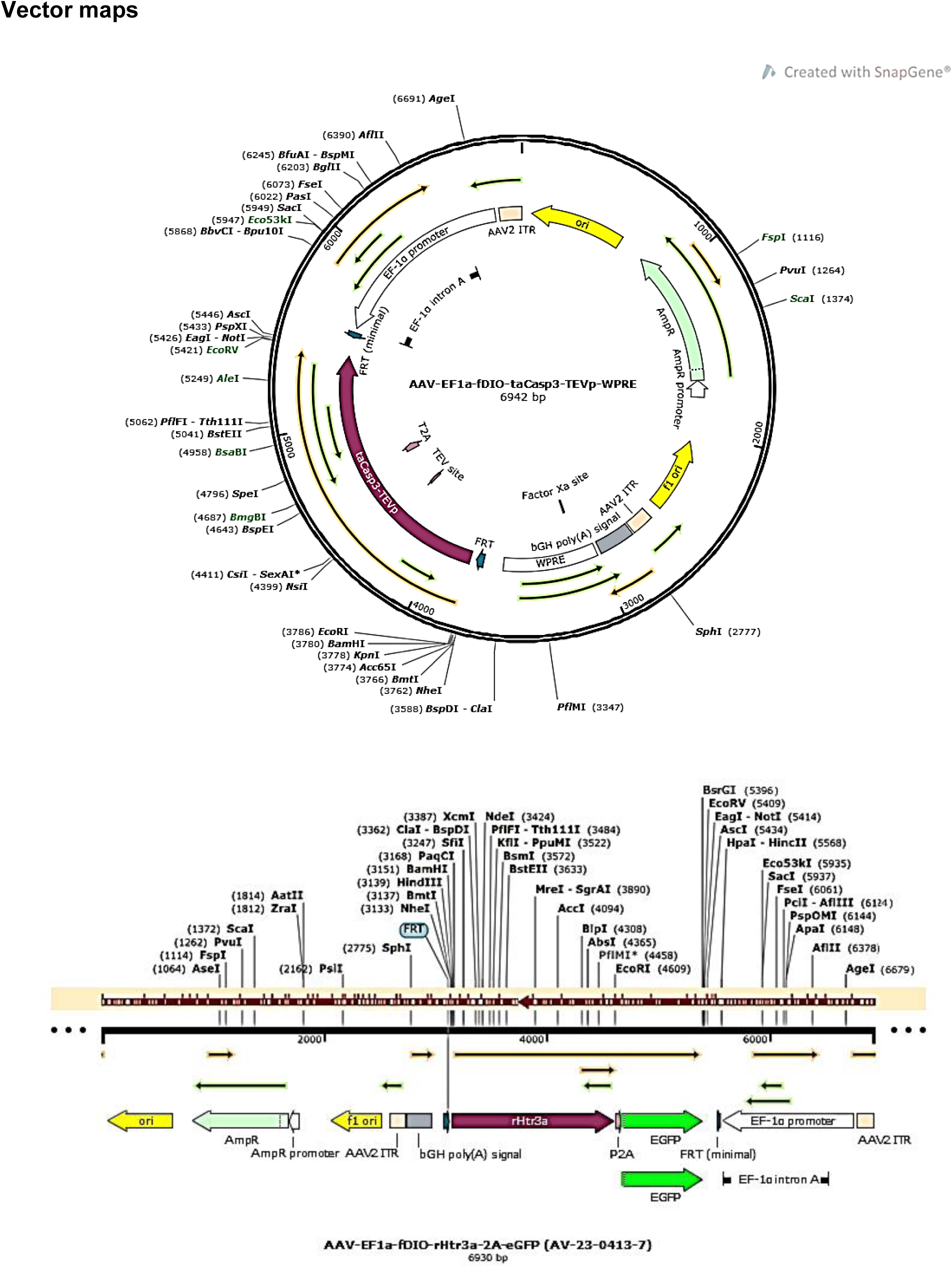

**Table 2.** List of antibodies.

| **Antibody** | **Host** | **Vendor** | **Catalogue Number** | **Type** |
| --- | --- | --- | --- | --- |
| GFP | Chicken | Aves Labs | GFP-1020 | Primary |
| Anti-Chicken IgY (H+L) Alexa Fluor™ 488 | Goat | Invitrogen | A-11039 | Secondary |

**Table 3.** Trp diet full content.

| **Product #** | **A20012701** |  |
| --- | --- | --- |
|  | **5% Trp** |  |
|  | gm% | kcal% |
| Protein | 22% | 23% |
| Carbohydrate | 64% | 66% |
| Fat | 5% | 12% |
| **Total** |  | **100%** |
| kcal/gm |  |  |
| **Ingredient (gm)** | gm | kcal |
| L-Arginine | 10 | 40 |
| L-Histidine-HCl-H<sub>2</sub>O | 6 | 24 |
| L-Isoleucine | 8 | 32 |
| L-Leucine | 12 | 48 |
| L-Lysine-HCl | 14 | 56 |
| L-Methionine | 6 | 24 |
| L-Phenylalanine | 8 | 32 |
| L-Threonine | 8 | 32 |
| **L-Tryptophan** | **50** | **200** |
| L-Valine | 8 | 32 |
| L-Alanine | 10 | 40 |
| L-Asparagine-H<sub>2</sub>O | 5 | 20 |
| L-Aspartate | 10 | 40 |
| L-Cystine | 4 | 16 |
| L-Glutamic Acid | 30 | 120 |
| L-Glutamine | 5 | 20 |
| Glycine | 10 | 40 |
| L-Proline | 5 | 20 |
| L-Serine | 5 | 20 |
| L-Tyrosine | 4 | 16 |
| **Corn Starch** | **502.5** | **2010** |
| Maltodextrin 10 | 125 | 500 |
| Cellulose | 50 | 0 |
| Corn Oil | 50 | 450 |
| Mineral Mix S10001 | 35 | 0 |
| Sodium Bicarbonate | 7.5 | 0 |
| Vitamin Mix V10001 | 10 | 40 |
| Choline Bitartrate | 2 | 0 |
| Yellow Dye, FD&C #5 | 0.05 | 0 |
| Red Dye, FD&C #40 | 0 | 0 |
| Blue Dye, FD&C #1 | 0 | 0 |
| **Total** | **1000.05** | **3872** |
| % Tryptophan | 5.00 |  |

**Extended Data Fig. 1:**
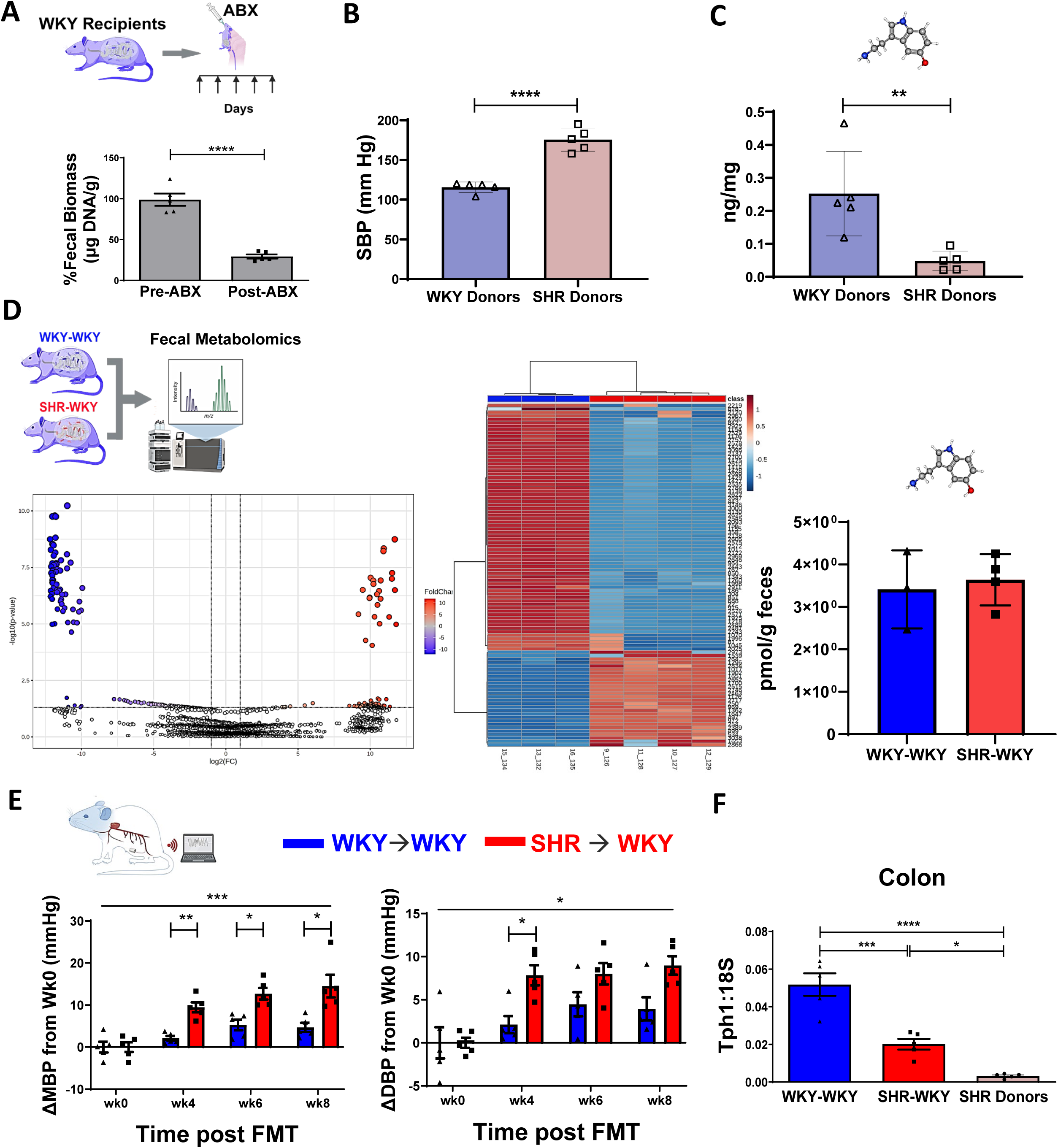
Hypertensive microbiota increased blood pressure and reduced colonic tryptophan hydroxylase 1 expression with no change in bacterial serotonin. **A,** five days of oral gavage of a cocktail of antibiotics (ABX) in Wistar-Kyoto (WKY) rats significantly reduced fecal bacterial DNA mass (by∼75%) confirming endogenous bacterial depletion. Values are means ± SEM; N=5/group; Student’s Ttest, ****P<0.0001. **B**, spontaneously hypertensive rats (SHR, light red) microbiota donors had significantly higher systolic blood pressures (SBP) than the Wistar-Kyoto (WKY, light blue) microbiota donors. Values are means ± SEM; N=5/group; Student’s Ttest, ****P<0.0001. **C**, SHR (light red) microbiota donors have significantly lower colonic serotonin levels (by∼2.5fold) than the Wistar-Kyoto (WKY, light blue) microbiota donors. Values are means ± SEM; N=5/group; Student’s Ttest, **P<0.01. **D**, top left, cartoon schematics of endpoint untargeted metabolomics performed in fecal bacterial samples of WKY→WKY (blue) and SHR→WKY (red) groups using LC-MS. The panel below and the middle panel reflect significant divergence of overall metabolites between the WKY→WKY (blue) and SHR→WKY (red) reflected in the volcano plot (bottom left) and heat map (middle panel) (N=4-5/group). Far right panel illustrates no significant difference in serotonin (5- HT) in fecal bacterial samples of WKY→WKY (blue) and SHR→WKY (red) group (N=5/group). Values are means ± SEM; N=4-5/group, Student’s Ttest. **E**, top left carton schematics illustrates radiotelemetric measurements of continuous blood pressures (BP, mmHg) performed once a week over 24hrs in conscious unrestrained WKY→WKY (blue) and SHR→WKY (red) rats at baseline (i.e., before fecal microbiota transplant (FMT), noted as wk0) and then at weeks 4-8. A significant increase in mean BP (MBP, by∼10mmHg) was observed in the SHR→WKY group (red) compared to WKY→WKY controls (blue) starting at wk4 post FMT (N=5/group). A significant increase in diastolic BP (DBP, by ∼6mmHg) was observed in the SHR→WKY group (red) compared to WKY→WKY controls (blue) at wk4 post FMT (N=5/group). Line across all time points reflects significant interaction between treatment and time for MBP and DBP. Weekly absolute values are normalized to values observed at baseline (wk0) for each rat and presented as change (Δ) from wk0. Values are means ± SEM; ANOVA with Mann-Whitney post-hoc, *P<0.05, **P<0.01, ***P<0.001. **F**, quantitative real time PCR (RT-qPCR) analysis for relative expression levels of tryptophan hydroxylase 1 (Tph1) normalized to 18S housekeeping gene in the proximal colons of the recipient groups and compared to SHR. A significant reduction in relative expression levels of Tph1 was observed in the colons of SHR→WKY rats (red) compared to the WKY→WKY controls (blue), reflecting a similar trend observed in the SHR. Values are means ± SEM; N=5/group; ANOVA with Mann-Whitney post-hoc, *P<0.05, ***P<0.001.

**Extended data Fig. 2:**
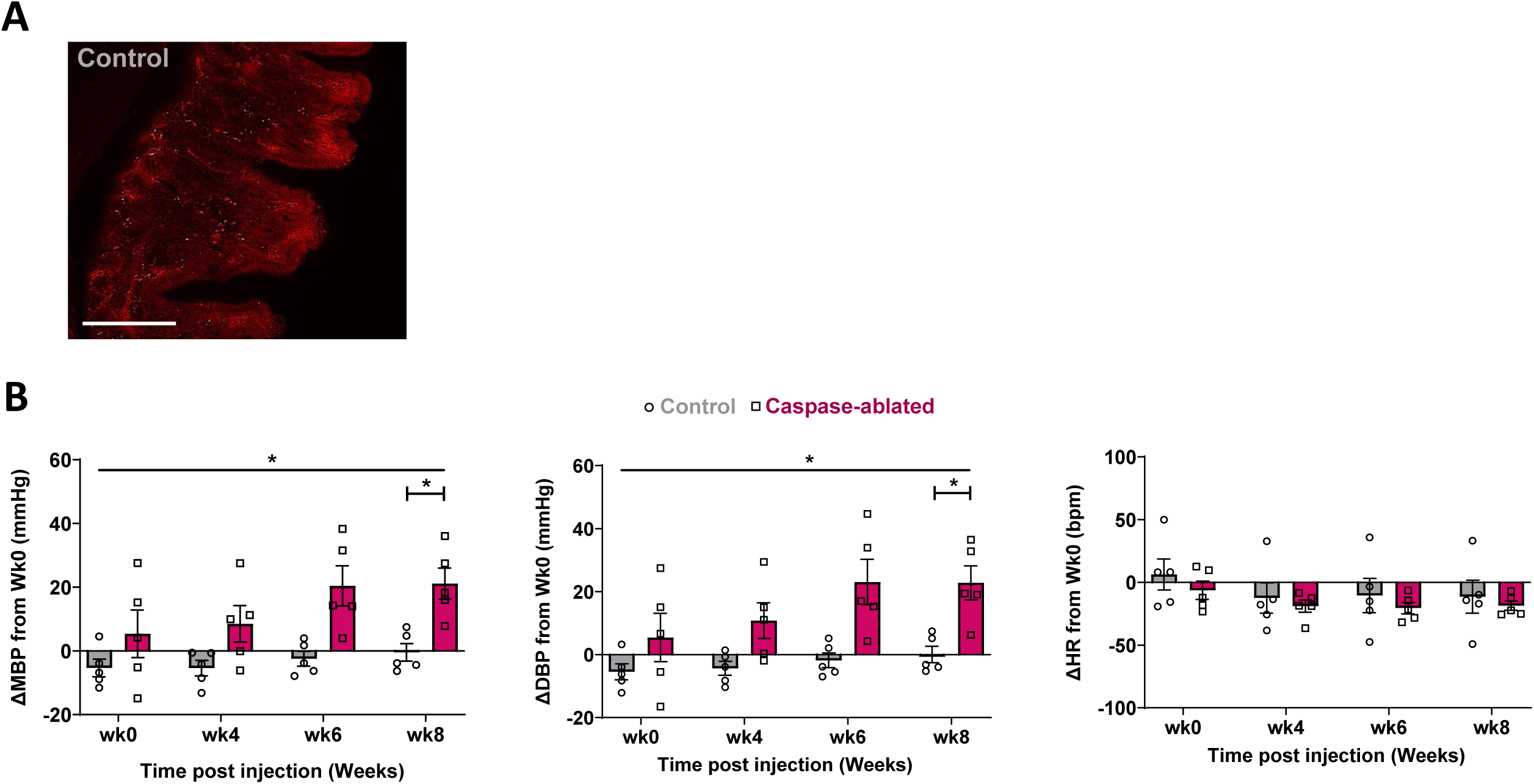
Chronic ablation of vagal afferents expressing 5HT3a receptors increases mean and diastolic blood pressure. **A,** proximal colons from Control--injected 5HT3aRCre rats were sectioned and imaged using fluorescence microscopy. tdTomato fluorescent labeling of vagal afferent projections was observed in proximal colons of Control-injected rats. Scale bar =100 µm. **B,** a significant increase in mean blood pressure (MBP, by∼20mmHg, left panel) and diastolic blood pressure (DBP, by ∼20mmHg, middle panel) was observed in 5HT3aR^Cre^ rats at wk8 following bilateral nodose ganglia microinjection of viral vector expressing Caspase 3 (Caspase-ablated, magenta) compared to the Control group (gray) (N=5/group). Line across all time points reflects significant interaction between treatment and time for MBP and DBP. No significant difference was observed in HR (right panel) between the groups at any time point. Absolute values are normalized to values observed at baseline for each rat and presented as change (Δ) from baseline. Values are means ± SEM; ANOVA with Mann-Whitney post-hoc, *P<0.05.

**Extended data Fig. 3:**
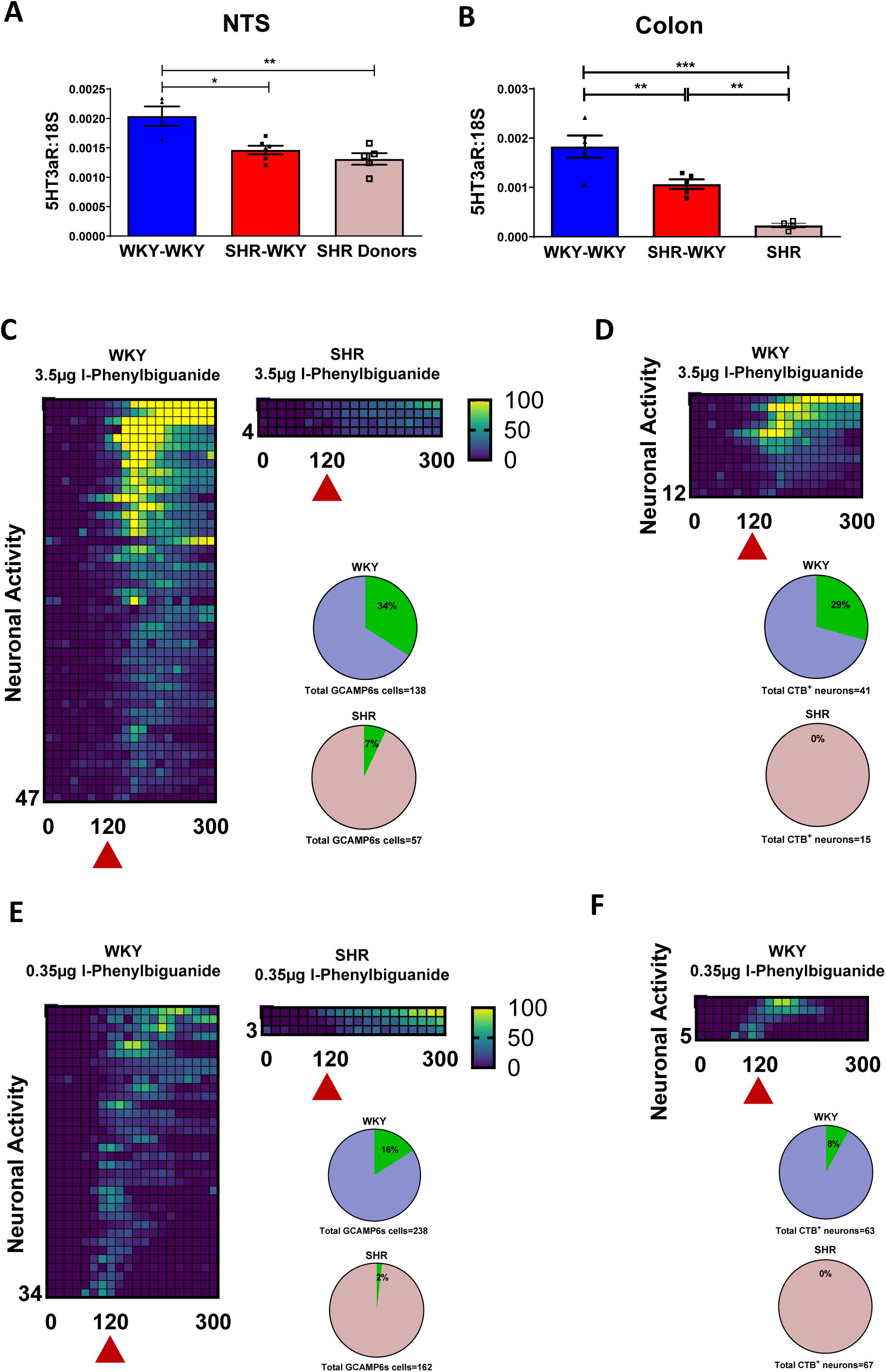
Reduced serotonin signaling by intestinal vagal afferents in hypertensive rodents with gut dysbiosis. **A-B,** quantitative real time PCR (qRT-PCR) analysis of relative expression levels of serotonergic 5HT3a receptors (5HT3aR) normalized to 18S housekeeping gene in the nucleus of the solitary tract (NTS, in A) and proximal colon (in B). A significant reduction in relative expression levels of 5HT3aRs was observed in the NTS and colon of SHR→WKY group (red) compared to WKY→WKY controls (blue) trending towards that observed in the SHR donors. Values are means ± SEM; N=5/group; ANOVA with Mann-Whitney post-hoc, *P<0.05, **P<0.01, ***P<0.001. **C,** heat maps depict z-score-normalized fluorescence traces from all GCaMP6s vagal neurons in the nodose ganglia of normotensive Wistar-Kyoto (WKY) rats (left panel) and spontaneously hypertensive rats (SHR, right panel), responding to i.v. infusion of serotonin receptor agonist L-phenylbiguanide (3.5µg). Each row represents the activity of a single cell over 5 mins. Stimulus was given at 120 seconds (red triangle). In N=5 per group. 34% of the total of 138 GCaMP6s-expressing neurons responded to the serotonin receptor agonist in the WKY rats, with only 7% of 57 GCaMP6s-expressing neurons were responsive in the SHR, as depicted by pie charts. The responses of each neuron are represented by heat maps. **D**, heat maps depict z-score-normalized fluorescence traces from all colon-projecting GCaMP6s vagal neurons in the nodose ganglia of normotensive Wistar-Kyoto (WKY) rats responding to i.v. infusion of serotonin receptor agonist (3.5µg). Each row represents the activity of a single cell over 5 mins. Stimulus was given at 120 seconds (red triangle). In N=5 per group, 29% of the total of the 41 colon-projecting GCaMP6s vagal neurons responded to the serotonin receptor agonist in the WKY rats, while none of the 15 colon-projecting GCaMP6s neurons were responsive in the SHR, as depicted by pie charts. The responses of each neuron are represented by heat maps. **E,** heat maps depict z-score-normalized fluorescence traces from all GCaMP6 vagal neurons in the nodose ganglia of normotensive Wistar-Kyoto (WKY) rats (left panel) and spontaneously hypertensive rats (SHR, right panel), responding to i.v. infusion of serotonin receptor agonist (0.35µg). Each row represents the activity of a single cell over 5 mins. Stimulus was given at 120 seconds (red triangle). In N=5 per group. 16% of the total of 238 GCaMP6s vagal neurons responded to the serotonin receptor agonist in the WKY rats, while only 2% of 162 GCaMP6s vagal neurons were responsive in the SHR, as depicted by pie charts. The responses of each neuron are represented by heat maps. **F**, heat maps depict z-score-normalized fluorescence traces from all colon-projecting GCaMP6s vagal neurons in the nodose ganglia of normotensive Wistar-Kyoto (WKY) rats (left panel) responding to i.v. infusion of serotonin receptor agonist (0.35µg). Each row represents the activity of a single cell over 5 mins. Stimulus was given at 120 seconds (red triangle). In N=5 per group, 8% of the total of the 63 colon-projecting GCaMP6s vagal neurons responded to the serotonin receptor agonist in the WKY rats, while none of the 67 colon-projecting GCaMP6s vagal neurons were responsive in the SHR, as depicted by pie charts. The responses of each neuron are represented by heat maps.

**Extended Data Fig. 4:**
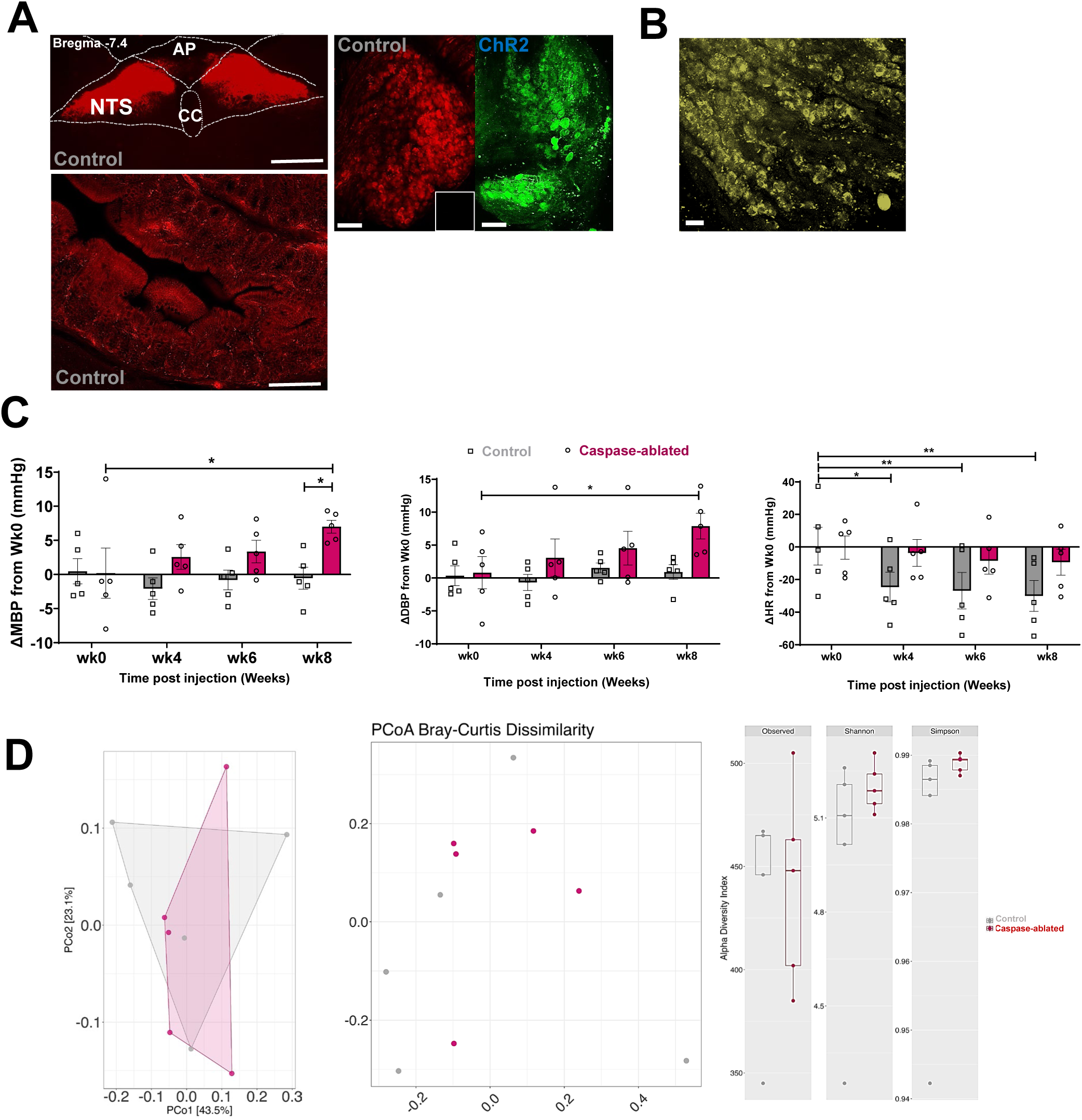
Viral ablation of colonic serotonin-sensing vagal neurons elevates blood pressure. **A,** endpoint confirmation of viral expression of fluorescence reporters in the nucleus of the solitary tract (NTS, top panel, scale bar = 300 µm), proximal colon (bottom panel, scale bar = 300 µm), and in the nodose ganglia of Control (left, with the inset image showing no green fluorescence in Control NG) and ChR2 (right, scale bar=100µm) in 5HT3aR^Cre^ rats injected with retrograde AAVrg in the proximal colon. AP-area postrema; CC-central canal. **B,** a representative image of viral expression of fluorescent reporter in colon-projecting vagal afferents in the nodose ganglia. The images are representative of viral transfection in the Control YFP group only, as the Caspase AAV construct does not express a fluorescent reporter (see details on viral constructs in Extended Data). Scale bar = 75µm. **C,** radiotelemetric measurements of continuous mean BP (MBP, mmHg), diastolic BP (DBP, mmHg) and heart rate (HR, beats per minute, bpm) were performed once a week over 24hrs in conscious unrestrained 5HT3aR^Cre^ rats at baseline (i.e., before nodose ganglia viral injections, noted as wk0) and then at weeks 4-8. A significant increase in MBP (by∼8mmHg, left panel) was observed in the Caspase-ablated group (magenta) compared to Controls (gray) and to its own baseline (wk0) at wk8 post injection. In addition, a significant increase in DBP (by ∼8mmHg, middle panel) was observed in the Caspase-ablated group (magenta) at wk8 post-injection compared to its baseline (wk0). Lastly, a significant decrease in HR (by ∼20-25 bpm) was observed at all time points in the Control (gray) but not the Caspase-ablated group (magenta). Absolute values are normalized to values observed at baseline for each rat and presented as change (Δ) from baseline. N=5/group. Values are means ± SEM; ANOVA with Mann-Whitney post-hoc, *P<0.05. **D**, endpoint 16S bacterial sequencing analyses show an overlap in composition and abundances of the gut bacteria between the two groups, illustrated by the principal coordinate analysis (PCoA) showing beta diversity with Bray-Curtis Dissimilarity and Alpha diversity plots with the common indices (far right).

**Extended data Fig. 5.**
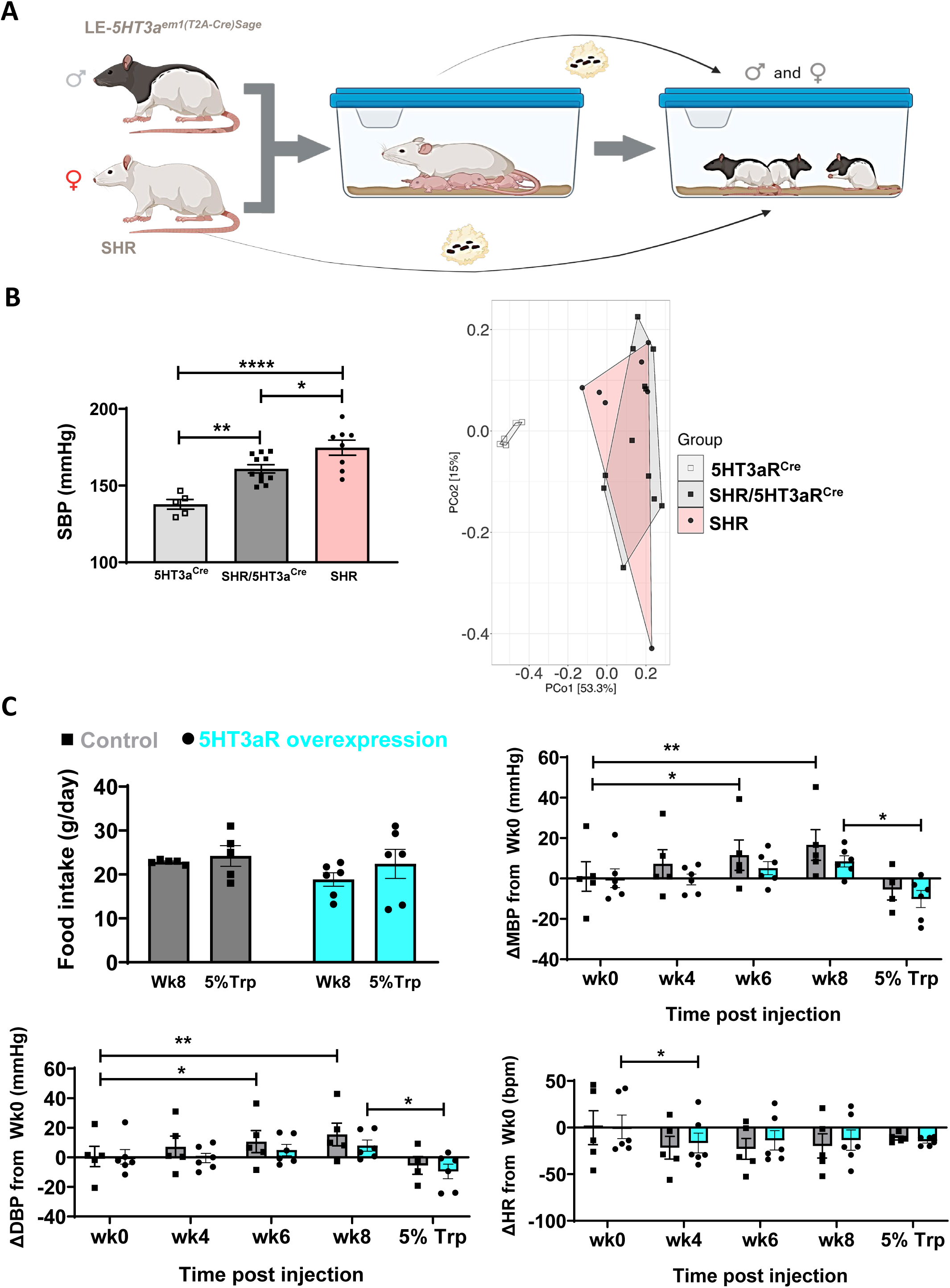
Viral expression of serotonin receptors in intestinal vagal afferents decreases blood pressure in hypertensive SHR/5HT3aR^Cre^ rats with gut dysbiosis. **A,** cartoon schematic depicting generation of the SHR/5HT3aR^Cre^ model. Female SHR were bred to male LE-5ht3a*^em1(T2A-Cre)Sage^* (5HT3aR^Cre^) rats. Following weaning, the offspring were transplanted with SHR gut microbiota in bedding, twice weekly until adulthood. **B**, adult SHR/5HT3aR^Cre^ rats presented with significantly higher systolic blood pressures (SBP, mmHg) compared to the 5HT3aR^Cre^ controls (left panel) and trending towards the SHR. On the right, principal coordinate analysis (PCoA) showing beta diversity following 16S sequencing of fecal bacteria revealed overlapping microbiota profiles between the SHR donors (light red) and the SHR/5HT3aR^Cre^ offspring recipients of the SHR microbiota (dark gray). In addition, a significant separation in the gut microbiota composition and abundances was demonstrated between the SHR/5HT3aR^Cre^ model (dark gray) and the 5HT3aR^Cre^ controls (light gray). **C**, top left shows no difference in the food intake before and during administration with 5% Trp in diet between the groups. Radiotelemetric measurements of continuous mean BP (MBP, mmHg), diastolic BP (DBP, mmHg) and heart rate (HR, beats per minute, bpm) were performed once a week over 24hrs in conscious unrestrained SHR/5HT3aR^Cre^ rats at baseline (i.e., before nodose ganglia viral injections noted as wk0) and then at weeks 4-9. Absolute values are normalized to values observed at baseline for each rat and presented as change (Δ) from baseline. N=4-6/group. Values are means ± SEM; ANOVA with Mann-Whitney post-hoc, *P<0.05, **P<0.01.

## Notes

### Competing Interest Statement

The authors have declared no competing interest.

### Summary of Updates

Grants and Funding section was added for clarification.

